# Genetics in the ocean’s twilight zone: Population structure of the Mueller’s pearlside across its distribution range

**DOI:** 10.1101/2025.02.22.639266

**Authors:** María Quintela, Alejandro Mateos-Rivera, Roger Lille-Langøy, François Besnier, Konstantinos Tsagarakis, Naiara Rodríguez-Ezpeleta, Miguel Bao, Martin Wiech, Lucilla Giulietti, Sonia Rábade-Uberos, Fran Saborido-Rey, Malika Chlaida, Espen Strand, Sofie Knutar, Eva García-Seoane, Webjørn Melle, Bjørn-Erik Axelsen, Kevin A. Glover

## Abstract

The large estimates of mesopelagic fish biomass have fuelled harvesting interests in the relatively untouched ocean’s twilight zone. The Mueller’s pearlside, one of the most abundant species inhabiting the north Atlantic mesopelagic layer, is a candidate to such fisheries despite its enormous ecological importance and the insufficient knowledge about its population genetic structure. To shed light on the latter, 863 individuals sampled across the North Atlantic and Mediterranean were genotyped using 170 genome-wide SNPs. Analyses revealed habitat-driven differentiation in three units: Mediterranean Sea, oceanic samples, and Norwegian fjords. These groups were not completely isolated to each other as a cline of Mediterranean admixture was detected in the Eastern Atlantic façade up to 47°N in an otherwise genetically homogeneous oceanic cluster. Temperature seemed to modulate the differentiation patterns, and in the Mediterranean added to the complicated topography of the Greek Seas to shape genetic structure. In the Norwegian coastline, sills did not hamper genetic exchange among fjords ranging 200 km apart, probably due to the position of the species in the water column together with its swimming capacity. This genetic information should be combined with demographic properties to outline the management of this species prior to any eventual fishery attempt.

## INTRODUCTION

The mesopelagic zone, located between 200–1000 m depth, and hosting up to 90% of the total oceanic fish biomass (Irigoien et al., 2014), is a largely unexplored area. The limited amount of sunlight at these depths explains the synonym ocean’s twilight zone, driving unique adaptations such as bioluminescence and diel vertical migration in the organisms inhabiting it. The enormous biomass of mesopelagic fish, ambitiously estimated around 10 Gt (10,000 million tonnes) (Irigoien et al., 2014), fuelled interests for the exploitation of a resource intended to aid meeting the demands of the expanding world population, both in terms of human food security and animal feed in a time where over a third of the worldwide fisheries operate beyond their biologically sustainable levels (FAO, 2020). However, a sustainable exploitation of the mesopelagic resources encounters challenges such as the limited knowledge about trophic interactions, life histories, behaviour, biomass, diversity, nutritional composition, and population genetic structure of the mesopelagic organisms (Hidalgo & Browman, 2019; Alvheim et al., 2020; Martin et al., 2020; Standal & Grimaldo, 2020; Wiech et al., 2020).

Mesopelagic organisms play a key ecological role in the Biological Carbon Pump (BCP), which is the suite of processes that collectively account for the ocean’s biologically driven sequestration of carbon from the atmosphere and land runoff to the ocean interior and seafloor sediments (Ducklow, 2001; Sigman & Haug, 2006; Boyd et al., 2019). Photosynthesis by phytoplankton in the sunlit surface ocean involves net uptake of CO_2_ from the atmosphere, whereas the transference of carbon from the surface to the deep ocean, amounting 1,300 Pg C yr^-1^, occurs via distinct pathways including gravitational settling of organic particles, mixing and advection of suspended organic carbon, as well as active transport by vertically migrating metazoans (Nowicki et al., 2022). The diel vertical migrations (DVM) of many mesopelagic fish species that feed on zooplankton near the surface at night and return to deep layers during the day actively export carbon from the surface to deeper water masses and contribute to the biological pump (Hidaka et al., 2001; Shreeve et al., 2009; Robinson et al., 2010) in what is regarded as the “largest daily migration of animals on earth” (Hays, 2003). The active transport mediated by diel vertical migration sequesters more carbon than the physical pump (subduction + vertical mixing of particles), *i.e.* 1.0 vs. 0.8 Pg C because of deeper remineralization depths (Stukel et al., 2023). Without it, the levels of atmospheric CO_2_ would be about 400 ppm higher than at present (Sanders et al., 2014; Boyd, 2015).

The Mueller’s pearlside, *Maurolicus muelleri* (Gmelin, 1789) is a small but abundant mesopelagic fish belonging to the family Sternoptychidae (see Grimaldo et al. (2020)). The genus *Maurolicus* was initially regarded as monotypic with a single species, *M. muelleri* (Gmelin, 1789), of cosmopolitan distribution. Later on, following identification of differences in combinations of morphometric characters, the genus was split into 15 species (Rees et al., 2020, see Fig. 1). However, many of them display overlapping ranges of meristic and morphometric traits and thus remain poorly characterized (Rees et al., 2017). Mitochondrial (16S and COI) and nuclear (ITS-2) gene sequences revealed groups conflicting with previously recognised species: (1) a ‘Northern’ clade comprising *M. muelleri* and *M. amethystinopunctatus*, (2) a ‘Southern’ clade comprising *M. australis*, *M. walvisensis* (also *M. japonicus*) and (3) an Eastern Equatorial and Western North Atlantic *M. weitzmani* (Rees et al., 2017). Synonymisation is proposed for *M. muelleri* and M*. amethystinopunctatus*, with limited morphological variation likely to reflect physical and biological differences experienced North/South of the sub-polar front (Rees et al., 2017). In a follow-up involving multiple locations worldwide, *M. muelleri* and *M. australis* were regarded as potentially a single species, but since no shared haplotypes were found between the two disjunct groups, they were kept separately (Rees et al., 2020). *M. muelleri* is thus distributed in the North Atlantic and Mediterranean Sea.

**Fig 1.**
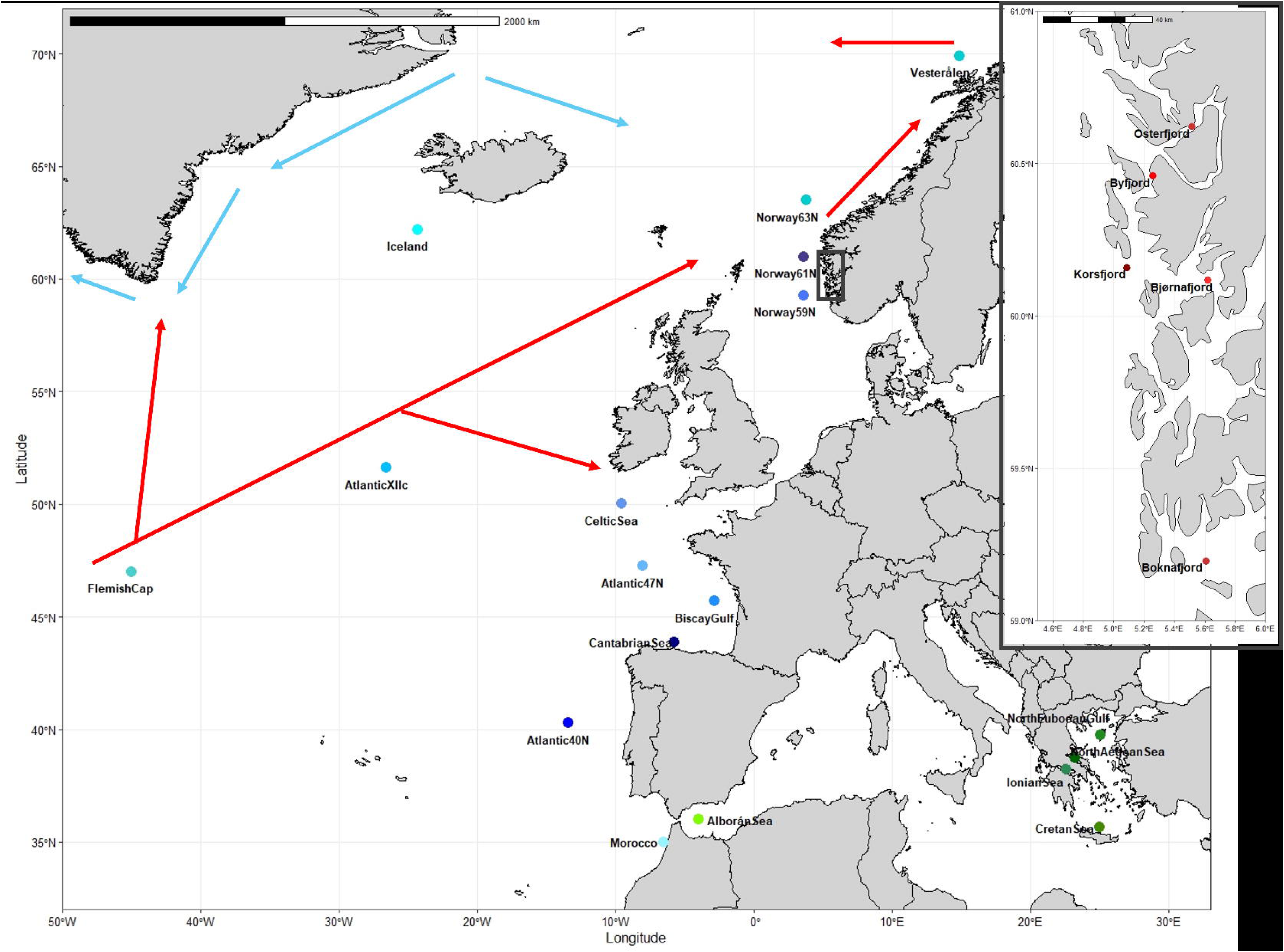
*Maurolicus muelleri* samples. Blue dots depict the Ocean samples whereas green dots depict the Mediterranean ones. The fjord samples, zoomed in the box to the right, are signalled with red dots. The main cold ocean currents are outlined in blue whereas the warm ones are outlined in red. For three of the sampling sites (*i.e.* Iceland, Cantabrian Sea and Gulf of Biscay) consisting of different nearby stations, the geographic coordinates correspond to the most central site.

*M. muelleri* is one of those species conducting DVM that play a critical role in marine ecosystems both as part of the BCP as well as linking primary consumers and higher tropic levels such as larger fish, cephalopods, seabirds and mammals (e.g. Cherel et al., 2010; Drazen & Sutton, 2017). The species forms distinct sound scattering layers at depth (SLs) and can be acoustically detected even from the youngest ontogenic stages due to their prominent swim bladder. In the Norwegian fjords, the SL composed of adults display DVM between April and June (Rasmussen & Giske, 1994; Goodson et al., 1995), whereas in early winter (January) adult fish remain in deeper waters while juveniles perform DVM as well as nocturnal midnight sinking after dusk (Giske et al., 1990; Baliño, 1993; Goodson et al., 1995). Fully grown individuals can achieve swimming speeds of 2–7 body lengths per second (i.e. 8–30 cm·s^-1^) (Torgersen & Kaartvedt, 2001).

In addition to taxonomic considerations, the literature dealing with population structure in *M. muelleri* is sparse and limited to few genetic studies. In western Norway, allozymes revealed that samples collected from five fjords displayed varying degrees of genetic differentiation to each other, and to a sample collected in the North Sea (Suneetha & Nævdal, 2001). In the Bay of Biscay, thousands of SNPs obtained from RADseq revealed no clear evidence of population structure as well as lower observed heterozygosity than anticipated (Rodriguez-Ezpeleta et al., 2017). Finally, mitochondrial DNA genes (COI, 12S, and 16S) revealed a lack of genetic structure among samples taken from the Greek Seas (Sarropoulou et al., 2022).

The correct outline of biologically correct management units or stocks is one of the multiple requirements of sustainable fisheries and intends to prevent the overexploitation of unique spawning components (e.g., see Kerr et al. (2017) for review). As this ecologically crucial species is a potential candidate for exploitation in a commercial fishery, there is a need to fill the knowledge gap regarding its ocean-wise genetic structure. In addition, when outlining management areas, special attention needs to be directed to identify locally adapted populations or subdivided ones as they might have different sustainable yield levels and be more prone to the negative effects of overfishing (Waples, Punt, and Cope 2008; Pinsky and Palumbi 2014). Furthermore, the range shifts that many marine species are already experiencing in the face of climate change should also be taken into consideration (Palacios-Abrantes et al. 2022; Dahms and Killen 2023).

The aim of the present study was to provide the first ocean-wide examination of genetic structure of *M. muelleri*. Several hundreds of individuals sampled across most of the species’ distribution range including both sides of the Atlantic (between longitudes 26 °W–25°E and latitudes 35–70 °N) were high-throughput genotyped using a suite of 170 SNP loci. Using the ecological parallelism with the mesopelagic glacier lanternfish *Benthosema glaciale* (Quintela et al., 2024), we hypothesize that the genetic patterns of differentiation of *M. muelleri* could be habitat-related with three major units corresponding to Norwegian fjords, open Atlantic ocean and Mediterranean, respectively.

## MATERIALS AND METHODS

### Sampling and genotyping

In the period 2017–2024, 941 individuals were collected in 23 sampling sites ranging across large part of the North Atlantic distribution of the species (Fig. 1). Based upon our previous experience with another widely distributed mesopelagic fish, *B. glaciale* (Quintela et al., 2024), three main habitats were aimed at: Norwegian fjords (*i.e.* five samples around 60 °N), Mediterranean Sea (*i.e.* one sample from the Alborán Sea plus four Eastern samples from the Greek Seas), and open sites in the Atlantic ocean (*i.e.* thirteen samples within latitudes 35 – 70 °N and longitudes 26 °W–25 °E). Wherever available, 50 individuals were collected per location. Two of the Greek samples came from the enclosed, deep and isolated gulfs of Corinth (Ionian Sea) and of North Euboea, with geophysical features similar to the Norwegian fjords (Kapelonis et al., 2023). Fin clips were taken and stored in ethanol 96% prior to DNA isolation, which was conducted using SPRI paramagnetic beads from the Beckman Coulter DNAdvance kit (A48706). DNA concentration was quantified using NanoDrop 8000.

Four sampling sites belonging to the different habitats were selected for SNP mining following the procedure used in Quintela et al. (2024): Mediterranean (Ionian Sea), fjords (Boknafjord) and oceanic (two sites in the Atlantic Ocean at latitudes 40 and 50 °N, *i.e.* Atlantic_40N and Celtic Sea, respectively). From these samples RNA-free DNA was extracted using the Beckman Coulter DNAdvance kit (A48706). DNA integrity was assessed by agarose gel electrophoresis and DNA concentration was quantified using Thermo Fisher Qubit dsDNA Broad Range (Q32853). Equimolar amounts of DNA from 10 individuals per site were pooled, and one library was prepared per pool using Illumina DNA prep. Pooled samples were then sequenced on a NovaSeq X using 1/8 of a NovaSeq X Series 25B flow cell (150 PE). The package *fastp* (Chen et al., 2018) was used for data preprocessing, which included deduplication, adaptor removal and analysis of over-represented sequences. FASTQC v0.11.5 (https://www.bioinformatics.babraham.ac.uk/projects/fastqc/) and MULTIQC v1.7 (https://github.com/MultiQC/MultiQC) were used to output graphics and statistics of the data quality control.

A draft assembly was produced using MEGAHIT (Li et al., 2015; Li et al., 2016) testing Kmer from 41 to 73. Scaffolding was performed with SoapDeNovo (https://github.com/aquaskyline/SOAPdenovo2) using 63 as Kmer value. The obtained draft assembly consisted mostly of small contigs, shorter than 500bp. However, 2156 contigs were larger than 10kb and represented together 27Mb. The four paired-end pooled samples were then mapped against this 27Mb short genome using BWA V0.7.17 (Li & Durbin, 2010) with default parameters for BWA aln and BWA sampe. Variant sites were called with the mpileup function (Li, 2011) from samtools V1.9 (Li et al., 2009). The R package *vcfR* (Knaus & Grünwald, 2016; Knaus & Grünwald, 2017) was used to visualise the distribution of coverage and mapping quality across all variant sites. Only Single Nucleotide Polymorphic (SNP) variants were retained if they fulfilled the following criteria: phred-scaled quality score (QUAL >500), coverage larger than 120X and lower than 300X for each pooled sample sites, and both alternative and reference allele represented in at least two samples out of four. This filtering produced 12,000 SNP that revealed significant differentiation among the three putative habitats as hypothesized. The differentiation was particularly large versus the Mediterranean according to the first axis of the PCA plot (PC1), which accounted for 39.5% of the variation (Suppl. Fig. S1) whereas PC2 (31.2%) depicted the differentiation between fjord and ocean as well as between oceanic samples. Finally, SNPs located less than 200 bases from another polymorphic site (SNP or indel) were removed to keep only SNPs with stable primer sequences. A total of 2034 SNPs fulfilled those criteria; however, the goal was to genotype as many individuals as possible across the species’ distribution range using a high-throughput SNP genotyping approach in a reasonably affordable manner. Therefore, the Assay Designer option of the Typer v5 program (Agena Biosciences CA, USA), was used to arrange the 2034 SNPs into different multiplex reactions with a maximum of 25 loci each. Based on number of SNPs and the multiplex confidence score, eight multiplexes containing in total 200 SNPs were finally selected to genotype the full set of samples (N=941). Primers were designed, and genotyped on the 941 individuals using the Sequenom MassARRAY iPLEX Platform as described by Gabriel et al. (2009). From the Flemish Cap (NW Atlantic), only 10 individuals were obtained, which were sampled both for fin clips and muscle. Thus, DNA was extracted from both tissues and individuals were amplified twice to conduct positive controls and reproducibility.

### Genetic structure

Statistical analyses were restricted to the subset of 170 well-functioning polymorphic loci obtained out of the 200 screened ones (Suppl. Table S1 compiles the reasons for discarding the 30 ones). Even though the threshold of acceptance of missing data per individual was 20%, only 3 of the 863 finally retained individuals showed the maximum missing data allowed, whereas 63% of them displayed ≤5%. These 863 individuals were distributed into 23 samples across three different habitats: fjords, Atlantic Ocean and Mediterranean Sea. To assess if the 170 SNPs would accurately discriminate between individuals in a population, the genotype accumulation curve was built using the function *genotype_curve* in the R (Team, 2020) package *poppr* (Kamvar et al., 2014) by randomly sampling x loci without replacement and counting the number of observed multilocus genotypes (MLGs). This repeated r times for 1 locus up to n-1 loci, creating n-1 distributions of observed MLGs. The observed (*H_o_*) and unbiased expected heterozygosity (*uH_e_*) as well as the inbreeding coefficient (*F*_IS_) were computed for each sample with GenAlEx v6.1 (Peakall & Smouse, 2006). Likewise, the genotype frequency of each locus and its direction (heterozygote deficit or excess) was compared with Hardy-Weinberg expectations (HWE) using the program GENEPOP 4.0.6 (Rousset, 2008) as was linkage disequilibrium (LD) between pairwise loci. The False Discovery Rate (FDR) correction of Benjamini and Hochberg (1995) was applied to *p*-values to control for Type I errors. Non-parametric Kruskal-Wallis’s rank sum test followed by post-hoc Dunn’s test were applied to perform comparison of estimates of genetic diversity among habitats. Data conversion into the appropriate formats was conducted using PGDSpider 2.1.1.5 (Lischer & Excoffier, 2012).

Unsupervised analysis of genetic structure were conducted with Principal Component Analysis (PCA) using the function *dudi.pca* in *ade4* package (Dray & Dufour, 2007) in R (Team, 2020) after replacing missing data with the mean allele frequencies, using no scaled allele frequencies (scale = FALSE). In addition, the Bayesian clustering approach implemented in STRUCTURE v.2.3.4 (Pritchard et al., 2000), and conducted using the software ParallelStructure (Besnier & Glover, 2013), was used to identify genetic groups under a model assuming admixture and correlated allele frequencies without using LOCPRIORS. Ten runs with a burn-in period consisting of 100,000 replications and a run length of 1,000,000 MCMC iterations were performed for K=1 to K=10 clusters. To determine the number of genetic groups, STRUCTURE output was analysed using two approaches: a) the *ad hoc* summary statistic ΔK of Evanno *et al*. (2005), and b) the Puechmaille (2016) four statistics (MedMedK, MedMeanK, MaxMedK and MaxMeanK), both implemented in StructureSelector (Li & Liu, 2018). Finally, the ten runs for the selected Ks were averaged with CLUMPP v.1.1.1 (Jakobsson & Rosenberg, 2007) using the FullSearch algorithm and the G’ pairwise matrix similarity statistic, and graphically displayed using bar plots.

Supervised genetic structure using geographically explicit samples was assessed using the Analysis of Molecular Variance (AMOVA) and pairwise *F*_ST_ (Weir & Cockerham, 1984), both computed with Arlequin v.3.5.1.2 (Excoffier et al., 2005). Furthermore, the relationship among samples was examined using the Discriminant Analysis of Principal Components (DAPC) (Jombart et al., 2010) implemented in the R (Team, 2020) package *adegenet* (Jombart, 2008) in which groups were defined using geographically explicit locations. To avoid overfitting, both the optimal number of principal components and discriminant functions to be retained were determined using the cross-validation function (Jombart & Collins, 2015; Miller et al., 2020).

The relationship between genetic (*F*_ST_) and geographic distance was examined to investigate if it adhered to the expectations of an “Isolation by Distance” pattern (IBD), *i.e.* increasing genetic differentiation with geographic distance as a result of restricted gene flow and drift (Wright, 1943; Slatkin, 1993; Rousset, 1997). A two-tailed Mantel (1967) test was conducted using PASSaGE v2 (Rosenberg & Anderson, 2011) and significance was assessed via 10,000 permutations. The matrix of pairwise shortest geographic distance by water (*i.e.* avoiding land masses) was calculated with the R (Team, 2020) package *marmap* (Pante & Simon-Bouhet, 2013).

Putative clines of allele frequency extending from the easternmost Atlantic coast into the Mediterranean were investigated via the latitudinal sliding-window approach developed by Pereira et al. (2018) using the R-scripts provided by the authors in Table S1.3. In a second step, loci identified as displaying clines were subjected to a geographic cline analysis conducted using the R package HZAR (Derryberry et al. 2014) over a circa 8000 km transect starting in Vesterålen and finishing in the Cretan Sea. The fifteen models implemented in HZAR were fitted to the allele frequency of the candidate loci to determine the position, width and shape of cline over the total geographic distance. The reference cline was built using STRUCTURE Q-score and, in both cases, the best cline model was decided upon AIC scores.

### Outlier detection and environmental association analyses

Loci departing from neutrality were identified using a combination of Arlequin v.3.5.1.2 (Excoffier et al., 2005) and BayeScan 2.1 (Foll & Gaggiotti, 2008). Analyses on Arlequin were conducted based on 100 demes with 50,000 simulations under a hierarchal island model. In BayeScan, sample size was set to 10,000 and the thinning interval to 50. Loci with a posterior probability over 0.99, corresponding to a Bayes Factor >2 (*i.e.,* “decisive selection” (Foll & Gaggiotti, 2006), were retained as outliers. The outcome of both methods was intersected to identify a panel of eventual consensus outliers.

Adaptation to local environments often occurs through natural selection acting on a large number of loci, each having a weak phenotypic effect. Temperature, salinity and dissolved oxygen are variables of physiological importance for marine ectotherms and of significance to account for myctophid distributions (e.g., Koubbi et al., 2011; Flynn & Marshall, 2013). Data on summer temperature measured at 200 m depth using a grid of ¼ ° and averaged for the period 2005-2012 was retrieved from NOAA database (National Oceanic and Atmospheric Administration). LFMM, “latent factor mixed model” (Frichot et al., 2013), was used to assess which variables could be a potential selective pressure driving local adaptation by identifying loci showing unusual associations with these environmental factors compared to the genetic background. This method uses a linear mixed model to test for associations between genetic variation and environmental factors, while controlling for neutral genetic structure with (random) latent factors. The number of latent factors was set at K=3 according to DAPC outcome and ten runs were conducted using 1,000 sweeps for burn-in and 10,000 additional sweeps. The corresponding z-scores of the ten replicates were combined following the recommendations described in Frichot and François (2015). First, the genomic inflation factor (λ) was obtained after computing the median of the squared (combined) z-scores for each K, divided by the median of the χ^2^ distribution with one degree of freedom. Finally, p-values were adjusted using the genomic inflation factor (λ) and false discovery rates were set using the Benjamini and Hochberg (1995) algorithm.

Redundancy Analysis (RDA), a genotype-environment association (GEA) method to detect loci under selection (Forester et al., 2018), was be used to analyse loci and several environmental predictors simultaneously. The SNP genotype matrix was used as the response matrix, while the environmental data matrix served as the explanatory variables. RDA determines how groups of loci covary in response to the multivariate environment, and can detect processes that result in weak, multilocus molecular signatures (Rellstab et al., 2015; Forester et al., 2018). Environmental data (temperature, salinity, pH, dissolved oxygen, current velocity, chlorophyll) was obtained for the different sampling points using the Bio-ORACLE database https://www.bio-oracle.org (Tyberghein et al., 2012; Assis et al., 2018). Collinearity between variable pairs was investigated and only non-correlated ones were retained for analyses. Analysis was conducted with the R package *vegan* v.2.5–7 (Oksanen et al., 2020). The candidate outliers obtained from the different analyses were annotated by matching the SNP flanking regions against non-redundant database of GenBank (www.ncbi.nlm.nih.gov/genbank/) using the Basic Local Alignment Search Tool (Altschul et al., 1990).

## RESULTS

### Genetic structure

SNPs genotyping of the 10 fish collected in the Flemish Cap which DNA was extracted in duplicate revealed similar performance for muscle and finclips, except in one of the individuals where finclips failed in 92% of the 200 tested loci. In the remaining nine pairs and excluding the differences due to failure of amplification in one of the individuals, 14 discrepancies (*i.e.* homozygote vs. heterozygote) were found (0.77%).

When considering the full set of individuals, the plateau of the genotype accumulation curve reached with 17 out of the 170 retained loci implied that 10% of the SNPs displayed enough power to discriminate between unique genotypes from the same sample (Suppl. Fig. S2). The Kruskal-Wallis test detected significant differences in genetic diversity across habitats (*P* of 0.005 and 0.002, respectively for *H*_O_ and u*H*_E_, and *P*=0.05 for the percentage of polymorphic markers) with lower values reported for the Mediterranean in all cases (Table 1). Dunn’s *a posterior* test revealed significant differences for the three estimates between Mediterranean and Oceanic habitats and between Fjords and Mediterranean for *H*_O_. Out of the 330395 performed tests for LD, 3% were significant after FDR. Deviations from HWE were found in 49 out of the 3910 loci by population tests after FDR (1.25%).

**Table 1.**
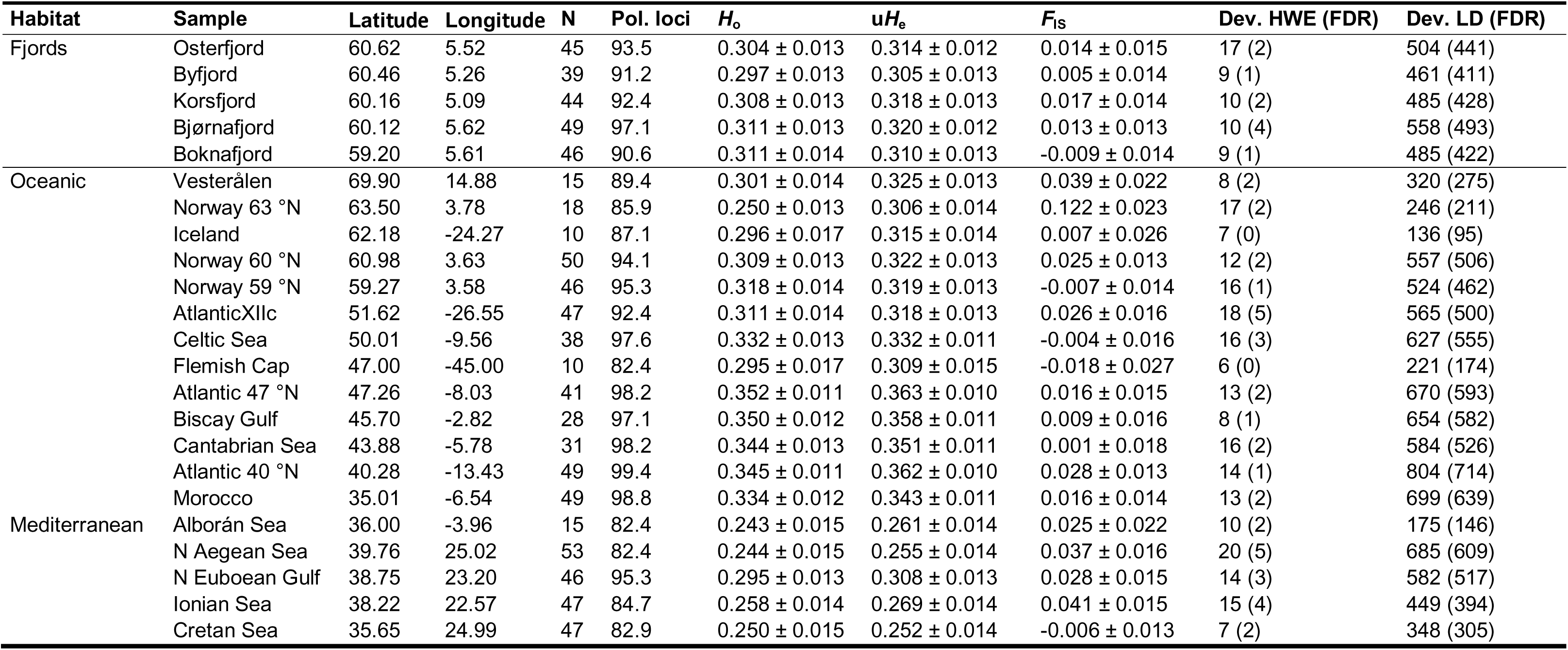
Sample summary statistics obtained for the set of 170 SNP loci: Sampling sites arranged into habitats with geographic coordinates in decimal degrees; number of individuals (N); percentage of polymorphic loci (Pol. loci); observed heterozygosity, *H*_o_ (mean ± SE); unbiased expected heterozygosity, u*H*_e_ (mean ± SE); inbreeding coefficient, *F*_IS_ (mean ± SE); number of deviations from Hardy-Weinberg equilibrium (Dev. HWE) and number of deviations from Linkage Disequilibrium (Dev. LD) at α=0.05 both before and (after) False Discovery Rate (FDR) correction. As some of the sampling sites consists of different nearby stations, the geographic coordinates indicated per sample are an average of all trawls for simplicity.

The first axis of the PCA biplot, accounting for 20% of the variation, neatly separated the Mediterranean from the fjords and part of the Oceanic individuals; however, a transition could be observed between the N. Euboean Gulf and some of the Atlantic samples such as Morocco, the Cantabrian Sea or the Gulf of Biscay (Fig. 2). The second axis (3.8%) separated the fjords and the Oceanic samples. In STRUCTURE *a posteriori* analyses, the Evanno test strongly suggested K=2 as the most likely number of genetic groups (ΔK=33400, Suppl. Fig. S3). This first division singled out the Mediterranean samples while revealing a decreasing cline of the Mediterranean genetic profile into the oceanic samples from Morocco northwards the Gulf of Biscay (Fig. 3a). The uniformity of the Mediterranean samples was disrupted in the Greek sample of the N. Euboean Gulf. At K=3, fjord and oceanic clusters became evident while the cline Mediterranean-Atlantic was still present in the southern Atlantic samples (Suppl. Fig. S4a). The Mediterranean sample from N. Euboean Gulf revealed a different cluster at K=5 (Suppl. Fig. S4c). Puechmaille’s solution of K=6 (Fig. 3b) reflected the genetic uniformity of the fjord samples, the distinctness and rather uniform Mediterranean genetic profile (excluding the N. Euboean Gulf) as well as two admixed clusters in the Atlantic Ocean (a northern one from Norway 63 °N to Flemish Cap and a southernmost one ranging from 47 °N to Morocco). Interestingly, from K=3 onwards, the sample from Vesterålen deviated from the oceanic profile and displayed a seemingly physical mixture of fish belonging to the fjord (majority) and northern Atlantic clusters.

**Fig 2.**
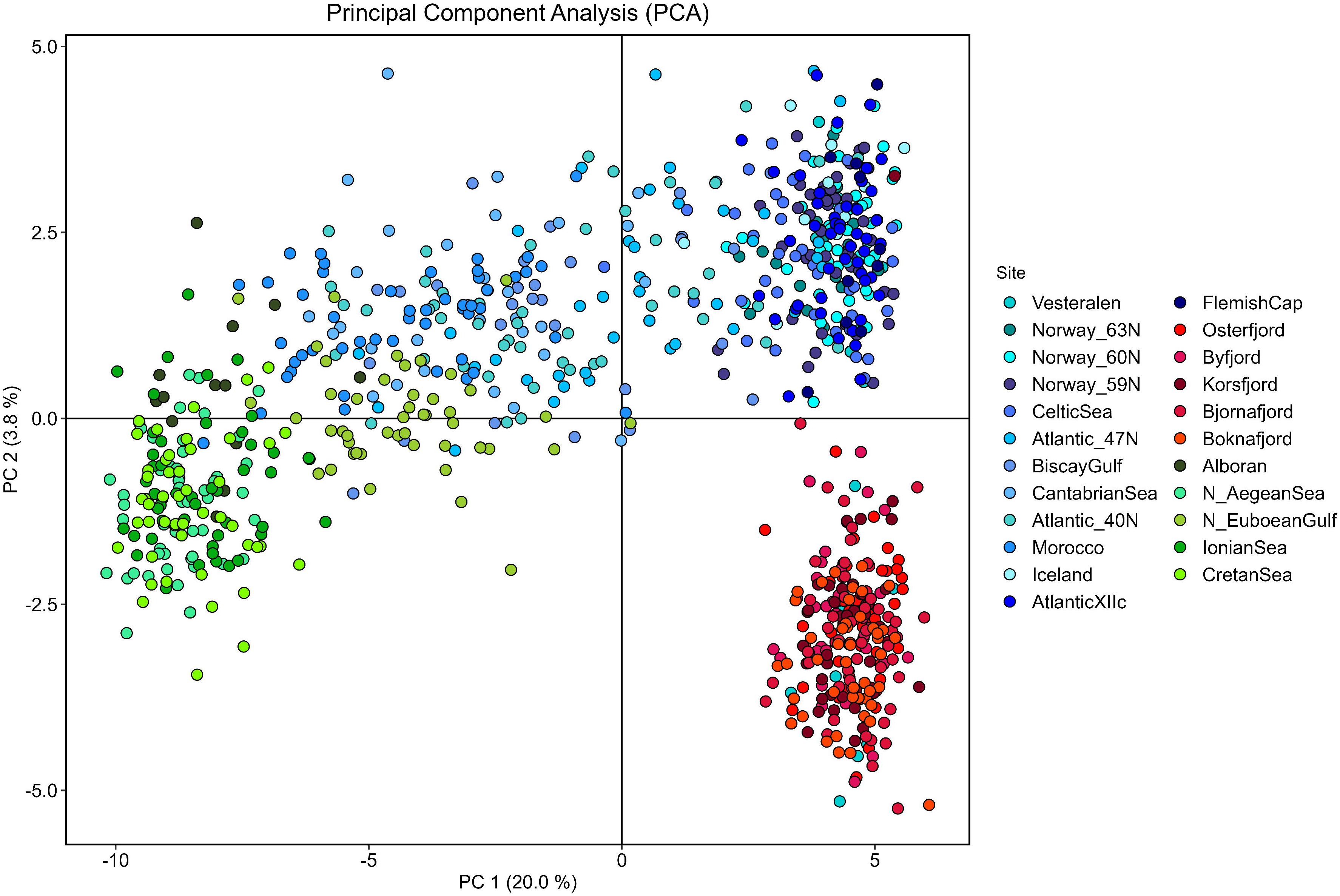
Principal Component Analysis (PCA) of *Maurolicus muelleri* genotyped at 170 loci. Blue dots depict the Ocean samples, red dots indicate fjord individuals and green dots represent the Mediterranean ones.

**Fig 3.**
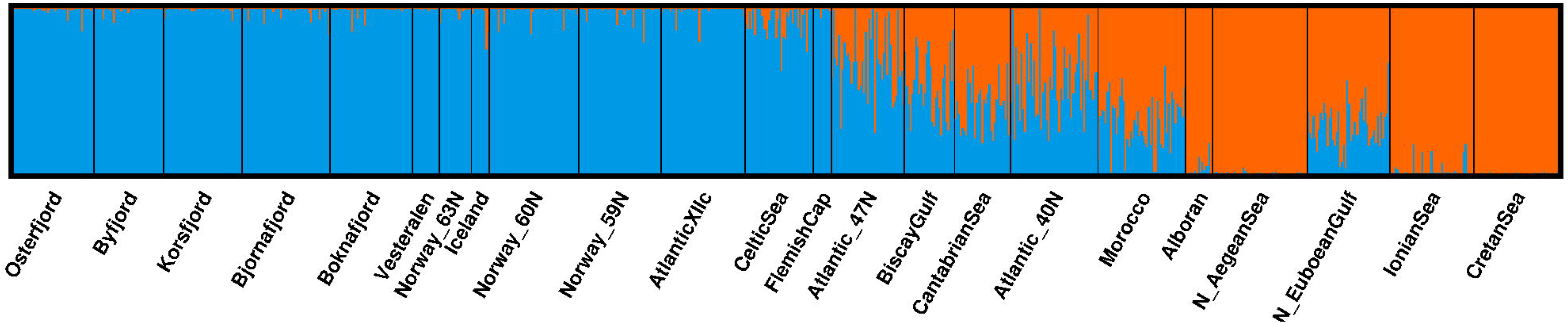
Barplot representing the proportion of individuals’ ancestry to cluster at a) K2 and b) K7 after Bayesian clustering in STRUCTURE as determined by the *a posteriori* analyses conducted using Evanno’s test and Puechmaille’s statistics, respectively.

High levels of genetic differentiation were detected on a per-locus basis, with 150 out of 170 loci significantly different from zero after FDR, and 83 of them with *F*_ST_ values ranging between 0.1 and 0.8. Hierarchical AMOVA revealed significant structure both among habitats (*F*_CT_=0.158, *P*<0.001), among samples within habitats (*F*_SC_=0.057, *P*<0.001) and within samples (*F*_ST_=0.206, *P*<0.001) with 15.8% of the variation hosted among habitats and 79.4% within samples. The dendrogram coupled with the pairwise *F*_ST_ (Fig. 4) revealed a first dichotomic division separating the Mediterranean, Morocco and the Cantabrian Sea from all the remaining sites. Interestingly, not only did the sample from the N. Euboean Gulf cluster with the latter Atlantic samples, but it also showed lower genetic differentiation towards both the Atlantic and the fjords than any of the Mediterranean samples (Suppl. Table S2). In the second branch of the tree, the fjords formed their own ramification while joining to Vesterålen whereas the Ocean samples were divided into the northernmost plus westernmost ones versus the ones between 47 and 40 °N. Most of the pairwise *F*_ST_ values were significantly different from zero (Suppl. Table S2) and strong differentiation was found among habitats, particularly towards the Mediterranean but exceptions were detected within habitats. The only significant differentiation found among the fjords occurred between the two farthest most sites, *i.e.* Osterfjord and Boknafjord separated by 200 km (*F*_ST_=0.007, *P*=0.001). In the Mediterranean, the only samples that did not differ from each other were the N. Aegean Sea and the Cretan Sea. The complicated topography of the Greek Seas with very confined body waters such as in the N. Euboean Gulf and the Gulf of Corinth (Ionian Sea) seems to hamper the genetic exchange whereas the 462 km of distance between the N. Aegean Sea and the Cretan Sea sampling sites did not seem to be a limitation for the gene flow. Within the oceanic samples, 71% of the pairwise comparisons were significant. However, almost no differentiation was detected between the samples ranging from Vesterålen southwards to the Celtic Sea together with the three westernmost ones (Iceland, AtlanticXIIc and FlemishCap). In the group of oceanic samples ranging between Morocco northwards to 47 °N, both the Cantabrian Sea and Morocco were significantly different from the remaining (pairwise *F*_ST_ ranging between 0.005 and 0.025) whereas no differentiation was detected among Atlantic_40 °N, Atlantic_47 °N and the Gulf of Biscay.

**Fig 4.**
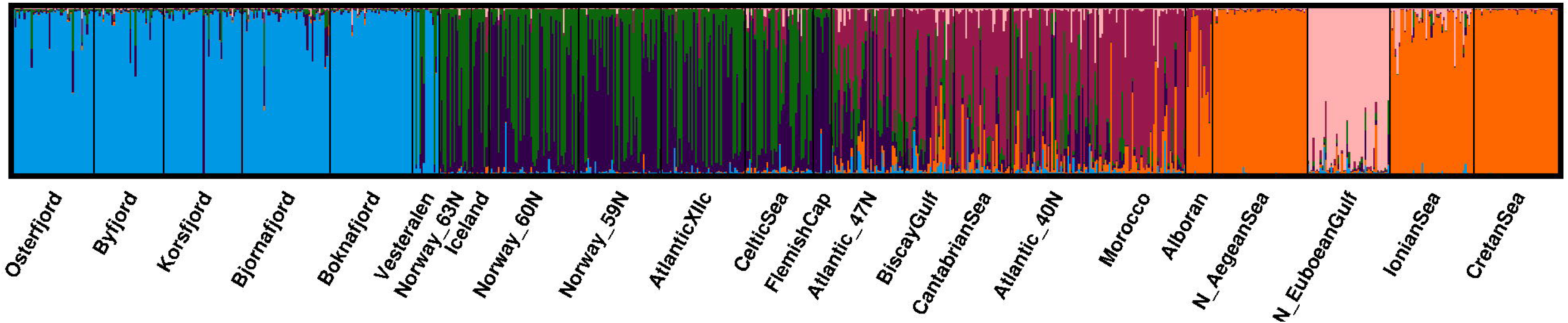
Heatmap *F*_ST_ coupled with dendrogram. Pairwise values and significance can be found in Suppl. Table S2. Colours depict the degree of differentiation: beige colours indicating lower differentiation and moving towards dark brown to indicate larger differentiation.

The DAPC revealed both differences among the three habitats (Ocean, Fjords and Mediterranean) as well as a certain continuity among them. Thus, the first axis of differentiation (60.8% of the variation) showed the uniqueness of the fjord individuals as well as a continuum between the Atlantic samples and the Mediterranean, together with the total lack of overlapping between the Mediterranean and the Northernmost and Westernmost Atlantic ones (Fig. 5a). The second axis of differentiation (16.8%) further singled out the fjord individuals. As formerly seen, some of the individuals from Vesterålen clustered with the fjord samples whereas the remaining clustered with the Northern group of the Atlantic. The third axis of differentiation (7%) singled out the sample from the N. Euboean Gulf (Fig. 5b). The positive and strong correlation (p=0.783, *P*<0.001) between the shortest water distance between samples and the genetic distance measured as pairwise *F*_ST_ was indicative of a pattern of Isolation by Distance (Fig. 6).

**Fig 5.**
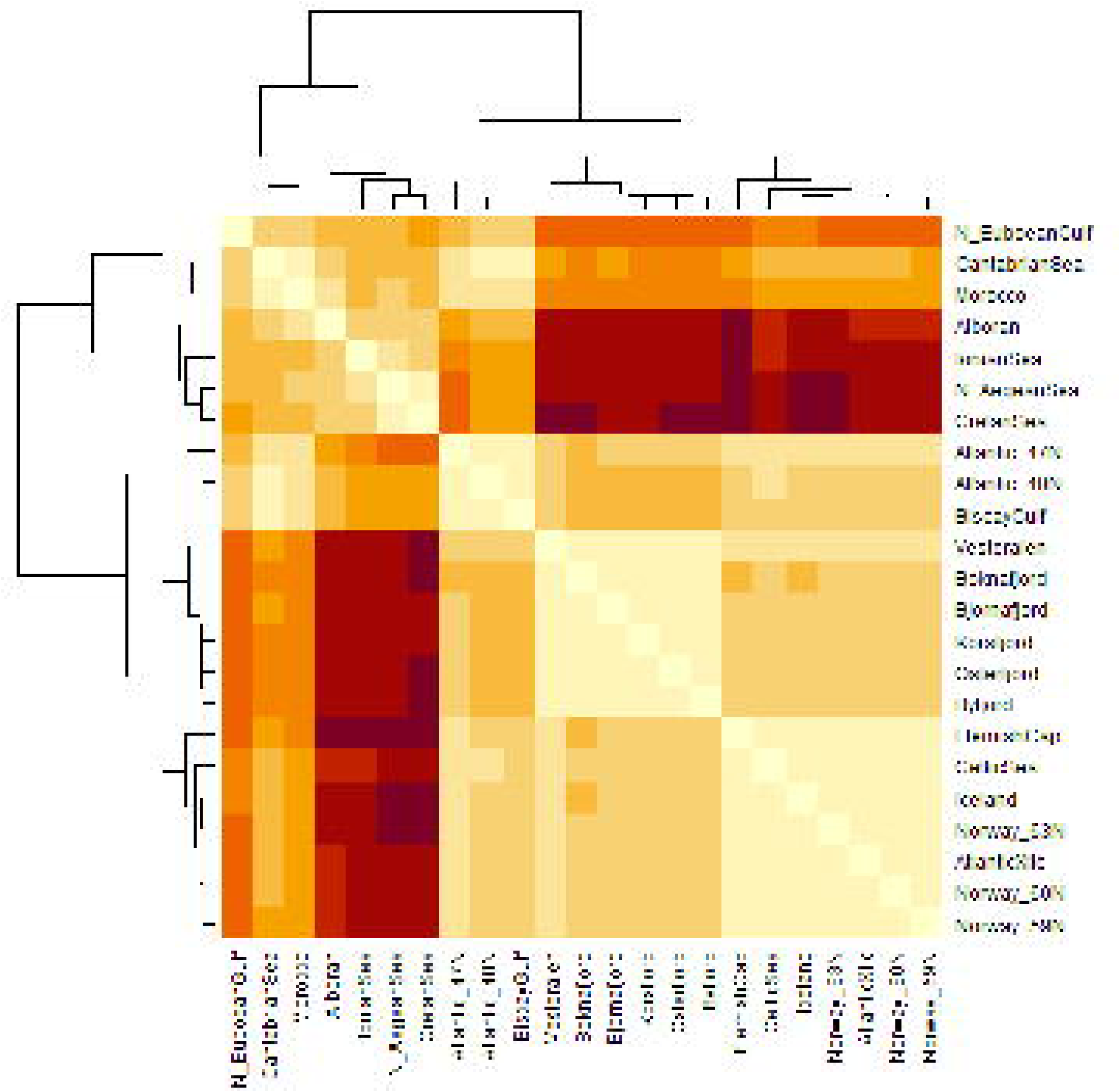
Genetic differentiation among *Maurolicus muelleri* samples assessed with 170 SNP loci using Discriminant Analysis of Principal Components (DAPC) after retaining 100 principal components and 3 discriminant functions: a) axis 1 and 2, and b) axis 1 and 3. Individuals from different sampling sites are represented by coloured dots.

**Fig 6.**
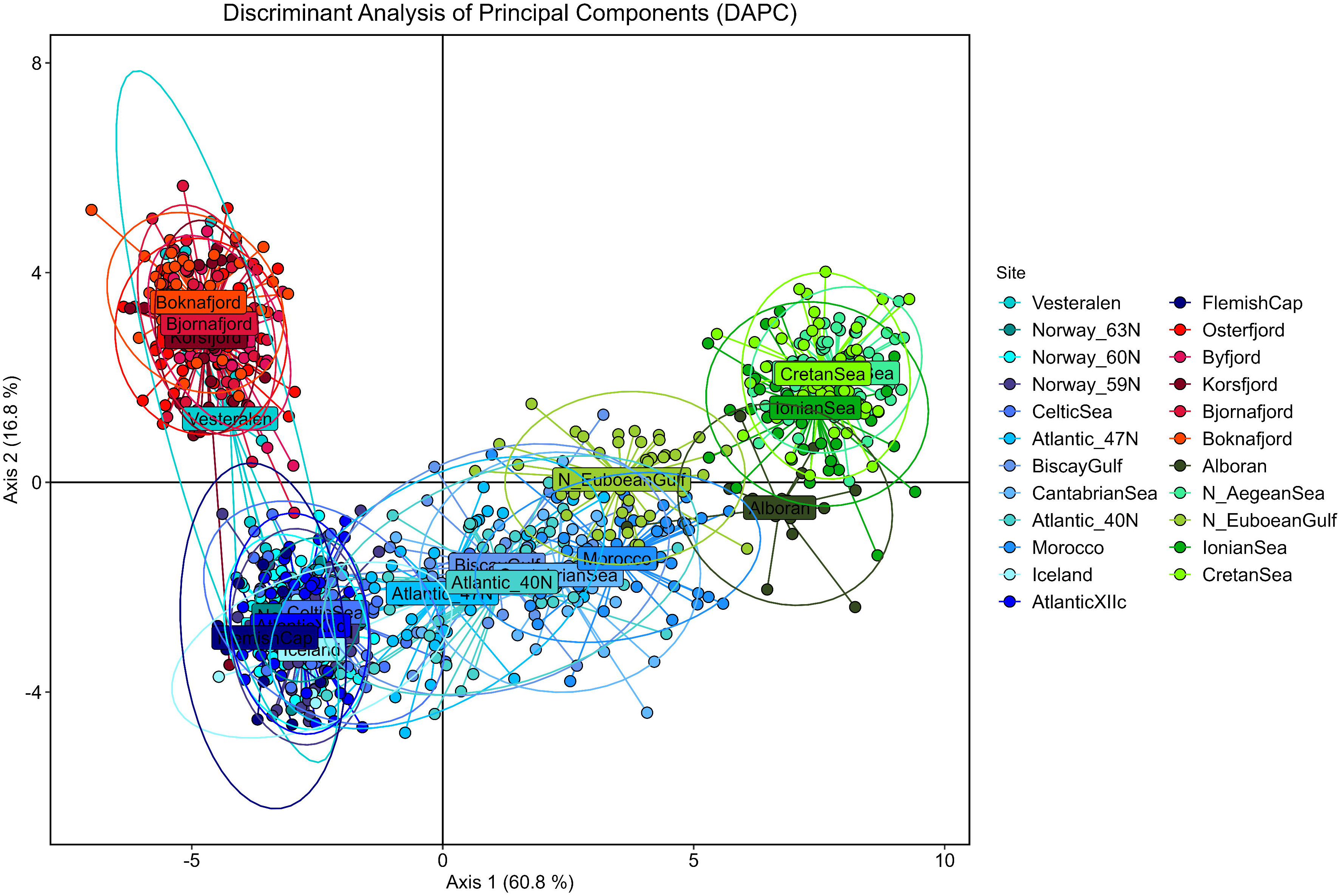
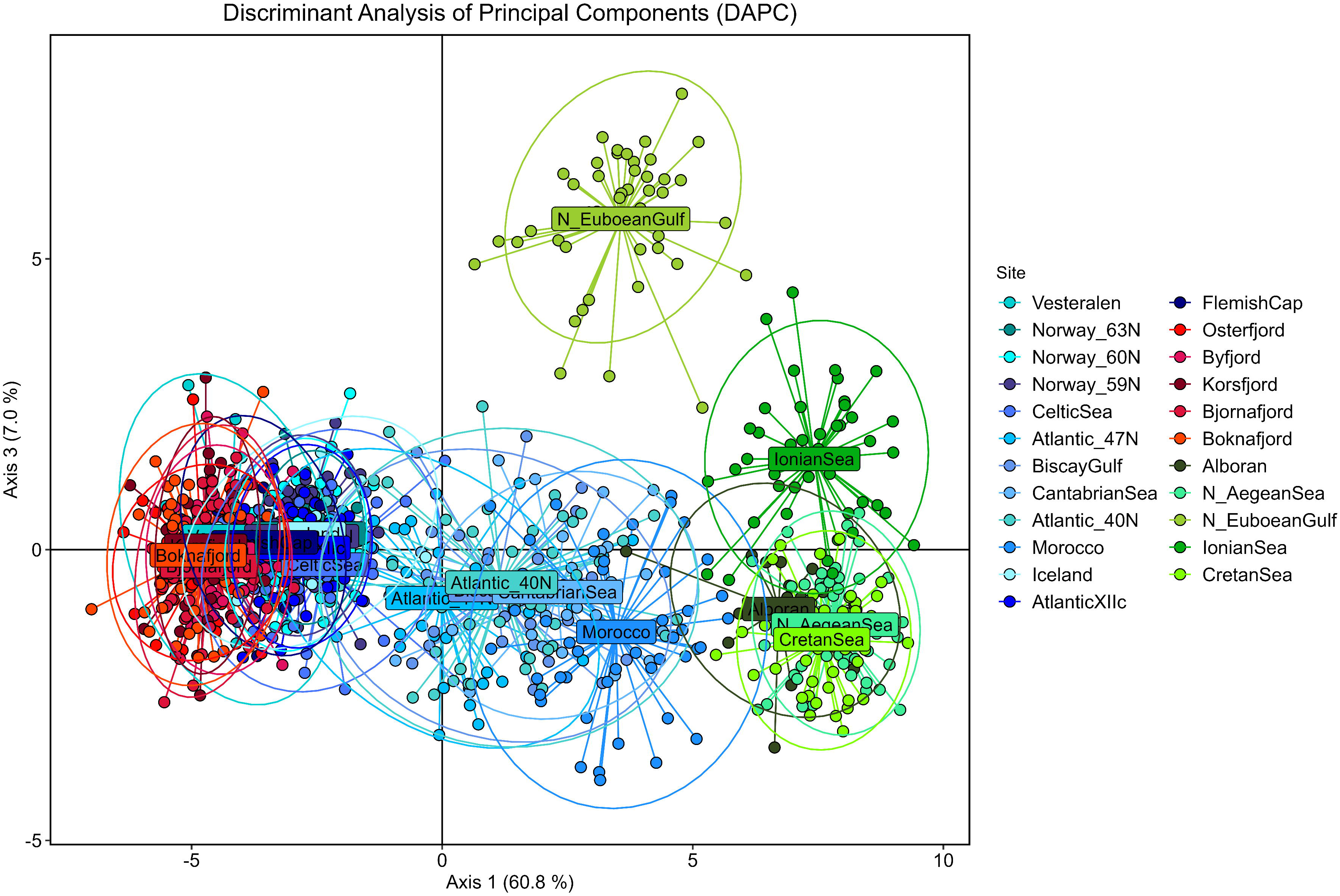
Mantel test showing significant correlation between shortest geographic distance by water (km) and genetic distance (*F*_ST_). Colours depict the samples involved in the comparisons within (red, blue, green) and across habitats (grey).

The allele frequencies of 72 out of the 170 loci showed signs of a latitudinal cline and only in one of them, the difference between maximum and minimum frequencies was <0.2 (in 49 of the loci, delta frequency ranged between 0.5 and 1). STRUCTURE Q-score also adhered to a clinal pattern with delta frequency of 0.99 (see Suppl. Fig. S5). HZAR revealed that for 18 of the 72 loci, the cline was centred within the limits of the reference cline, *i.e*. around 3000–3400 km south from Vesterålen (Fig. 7), which corresponds to a latitude around 44–47 °N.

**Fig 7.**
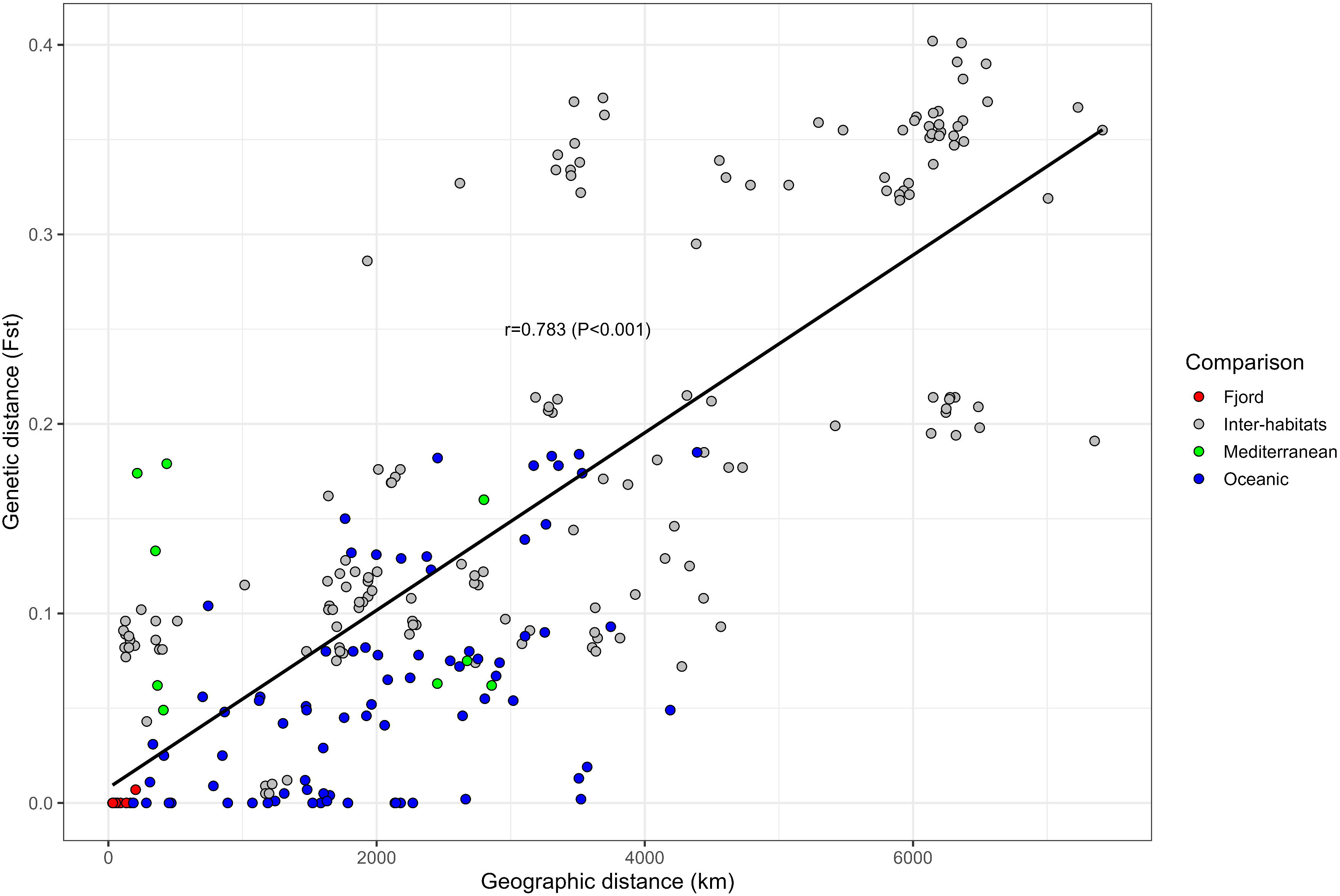
Summary of HZAR analyses: Cline centre and width for the loci showing signs of latitudinal clines. Red dotted lines represent the limits of the reference cline built with STRUCTURE Q-score.

### Outlier detection and environmental association analyses

Arlequin and BayeScan jointly flagged 20 loci as candidates to positive selection, 25 as putatively under balancing selection and 78 that did not depart from neutral expectations. The remaining 47 loci were found to be under selection only by one of the procedures. Data were divided into putatively identified neutral loci and loci under positive selection, and genetic structure was assessed separately.

In the neutral dataset, both the PCA biplot (Suppl. Fig. S6a) and the DAPC (Suppl. Fig. S6b) depicted the differentiation among habitats revealed by the set of total markers but with reduced capacity of discrimination of the transition between the Atlantic and the Mediterranean. Evanno test again suggested K=2 as the most likely number of genetic groups (ΔK=2105, Suppl. Fig. S7) and the resulting barplot mirrored the result obtained with the full set of loci (Suppl. Fig. S8a). The Puechmaille’s solution of K=6 (Suppl. Fig. S8b) revealed a similar pattern to the one with total loci without discriminating the Mediterranean sample of the N. Euboean Gulf. Hierarchical AMOVA revealed significant structure both among habitats (*F*_CT_=0.127, *P*<0.001), among samples within habitats (*F*_SC_=0.030, *P*<0.001) and within samples (*F*_ST_=0.154, *P*<0.001). The 12.7% of the variation was hosted among habitats whereas 84.6% was hosted within samples. Pairwise *F*_ST_ (Suppl. Fig. S9, Table S3) followed the patterns of the total loci.

The set of 20 candidate loci to positive selection lacked capacity of discriminating between fjord samples and oceanic ones, namely the Northern Atlantic cluster, both at PCA (Suppl. Fig. S10a) and the DAPC (Suppl. Fig. S10b). This lack of discrimination is illustrated by the three main groups of the *F*_ST_ dendrogram (Suppl. Fig. S11, Table S4): i) Norwegian fjords and Atlantic samples north of 47°, ii) Atlantic samples from 47 °N southwards plus the N_EuboeanGulf, iii) rest of Mediterranean samples.

LFMM analysis after adjusting p-values using the genomic inflation factor (λ) revealed that 21 loci were associated with summer temperature at 200 m depth. Of those loci, 19 were flagged as candidate outliers to positive selection both by BayeScan and Arlequin whereas the two remaining were only flagged by BayeScan. All loci identified by LFMM were involved in allele frequency clines (see Table 2 and Suppl. Table S5) with a break point at 47°N in the eastern part of the Atlantic. Interestingly, the allele frequency at locus P402 and P676 in N. Euboean Gulf fit better the frequency of the fjord and northern Atlantic samples than the Mediterranean ones, pattern that happens in some other loci and accounts for the distinctness of the N. Euboean sample and its relatedness to the Atlantic ones (see Fig. 3 or Fig. 5a). RDA identified two loci putatively related with temperature (P942 and P1458), both candidate outliers to positive selection, identified by LFMM as linked to temperature and with clines centered around 3230–3251 km. In addition, RDA identified a locus seemingly related to chlorophyll concentration (P452), which was flagged as under positive selection by BYSC and was not involved in any cline (Suppl. Table S5). BLAST identified two matches into zinc finger proteins, two in metalloproteinases, anoctamin and MAX interactor, respectively (Suppl. Table S5).

**Table 2.** LFMM analysis: Heatmap of allele frequency per sample for the candidate loci associated to summer average temperature measured from 2005 to 2012 at 200 m depth. Samples are distributed in blocks corresponding to habitats, *i.e.* fjords, oceanic and Mediterranean.

| Sample | Lat | Long | Temp (°C) | P105 | P253 | P359 | P402 | P635 | P676 | P709 | P942 | P989 | P1267 | P1446 | P1458 | P1500 | P1569 | P1606 | P1636 | P1691 | P1770 | P1856 | P1945 | P1989 |
| --- | --- | --- | --- | --- | --- | --- | --- | --- | --- | --- | --- | --- | --- | --- | --- | --- | --- | --- | --- | --- | --- | --- | --- | --- |
| Osterfjord | 60.62 | 5.52 | 7.49 | 1.000 | 0.773 | 0.789 | 1.000 | 0.769 | 1.000 | 1.000 | 1.000 | 0.975 | 0.989 | 0.956 | 0.989 | 0.956 | 1.000 | 0.878 | 1.000 | 0.878 | 0.778 | 0.978 | 1.000 | 0.977 |
| Byfjord | 60.46 | 5.26 | 7.49 | 1.000 | 0.686 | 0.763 | 1.000 | 0.809 | 1.000 | 1.000 | 1.000 | 1.000 | 1.000 | 0.974 | 1.000 | 0.962 | 1.000 | 0.949 | 1.000 | 0.816 | 0.692 | 0.974 | 1.000 | 0.962 |
| Korsfjord | 60.16 | 5.09 | 6.41 | 1.000 | 0.738 | 0.833 | 1.000 | 0.915 | 0.989 | 1.000 | 1.000 | 0.985 | 1.000 | 0.976 | 1.000 | 0.951 | 1.000 | 0.886 | 1.000 | 0.807 | 0.773 | 0.959 | 1.000 | 0.983 |
| Bjørnafjord | 60.12 | 5.62 | 6.41 | 0.990 | 0.719 | 0.745 | 1.000 | 0.902 | 0.990 | 0.989 | 0.990 | 0.978 | 0.969 | 0.959 | 0.990 | 0.949 | 0.989 | 0.957 | 1.000 | 0.786 | 0.796 | 0.959 | 1.000 | 0.990 |
| Boknafjord | 59.20 | 5.61 | 6.09 | 1.000 | 0.727 | 0.783 | 1.000 | 0.910 | 1.000 | 1.000 | 1.000 | 0.986 | 1.000 | 0.957 | 1.000 | 0.946 | 1.000 | 0.933 | 1.000 | 0.772 | 0.826 | 0.957 | 1.000 | 1.000 |
| Vesterålen | 69.90 | 14.88 | 6.04 | 1.000 | 0.786 | 0.800 | 1.000 | 0.900 | 1.000 | 1.000 | 1.000 | 0.955 | 1.000 | 0.955 | 1.000 | 1.000 | 1.000 | 0.900 | 1.000 | 0.833 | 0.800 | 1.000 | 1.000 | 1.000 |
| Norway_63N | 63.50 | 3.78 | 7.28 | 1.000 | 0.618 | 0.733 | 1.000 | 0.607 | 0.972 | 1.000 | 1.000 | 1.000 | 1.000 | 0.929 | 1.000 | 0.964 | 1.000 | 0.900 | 0.971 | 0.735 | 0.750 | 0.964 | 1.000 | 1.000 |
| Iceland | 62.18 | -24.27 | 8.40 | 0.950 | 0.650 | 0.444 | 0.950 | 0.500 | 1.000 | 0.950 | 0.950 | 0.950 | 0.950 | 0.944 | 0.950 | 0.944 | 0.950 | 0.800 | 0.950 | 0.750 | 0.833 | 0.944 | 1.000 | 1.000 |
| Norway_60N | 60.98 | 3.63 | 7.72 | 1.000 | 0.625 | 0.844 | 0.979 | 0.704 | 1.000 | 1.000 | 0.990 | 1.000 | 1.000 | 0.969 | 0.990 | 0.969 | 0.990 | 0.862 | 0.970 | 0.667 | 0.720 | 0.969 | 1.000 | 0.990 |
| Norway_59N | 59.27 | 3.58 | 7.61 | 0.989 | 0.543 | 0.731 | 0.978 | 0.674 | 0.989 | 0.989 | 1.000 | 0.950 | 0.989 | 0.977 | 1.000 | 0.976 | 1.000 | 0.891 | 0.967 | 0.682 | 0.824 | 0.975 | 0.989 | 0.987 |
| AtlanticXIIc | 51.62 | -26.55 | 10.44 | 1.000 | 0.717 | 0.696 | 1.000 | 0.649 | 1.000 | 0.989 | 0.968 | 0.966 | 0.989 | 1.000 | 0.968 | 1.000 | 0.968 | 0.943 | 1.000 | 0.707 | 0.733 | 1.000 | 1.000 | 0.967 |
| CelticSea | 50.01 | -9.56 | 10.90 | 0.932 | 0.595 | 0.724 | 0.905 | 0.618 | 0.956 | 0.908 | 0.908 | 0.909 | 0.908 | 0.935 | 0.908 | 0.952 | 0.919 | 0.842 | 0.947 | 0.618 | 0.816 | 0.938 | 0.961 | 0.938 |
| Flemish Cap | 47.00 | -45.00 | 4.38 | 1.000 | 0.650 | 0.900 | 1.000 | 0.650 | 0.944 | 1.000 | 1.000 | 0.850 | 1.000 | 1.000 | 1.000 | 1.000 | 1.000 | 0.938 | 1.000 | 0.556 | 0.950 | 1.000 | 1.000 | 1.000 |
| Atlantic_47N | 47.26 | -8.03 | 12.01 | 0.667 | 0.488 | 0.541 | 0.634 | 0.551 | 0.782 | 0.654 | 0.646 | 0.729 | 0.675 | 0.750 | 0.646 | 0.756 | 0.671 | 0.663 | 0.667 | 0.341 | 0.524 | 0.725 | 0.720 | 0.675 |
| Biscay Gulf | 45.70 | -2.82 | 12.08 | 0.593 | 0.389 | 0.463 | 0.571 | 0.214 | 0.654 | 0.589 | 0.611 | 0.769 | 0.589 | 0.455 | 0.611 | 0.458 | 0.574 | 0.673 | 0.411 | 0.407 | 0.346 | 0.435 | 0.571 | 0.563 |
| Cantabrian Sea | 43.88 | -5.78 | 11.62 | 0.233 | 0.345 | 0.293 | 0.597 | 0.226 | 0.536 | 0.250 | 0.419 | 0.714 | 0.250 | 0.281 | 0.419 | 0.342 | 0.433 | 0.450 | 0.371 | 0.210 | 0.355 | 0.313 | 0.500 | 0.525 |
| Atlantic_40N | 40.28 | -13.43 | 12.92 | 0.500 | 0.438 | 0.467 | 0.533 | 0.313 | 0.745 | 0.490 | 0.573 | 0.670 | 0.500 | 0.500 | 0.571 | 0.478 | 0.571 | 0.656 | 0.500 | 0.327 | 0.385 | 0.489 | 0.622 | 0.556 |
| Morocco | 35.01 | -6.54 | 14.19 | 0.208 | 0.240 | 0.260 | 0.234 | 0.202 | 0.500 | 0.189 | 0.306 | 0.511 | 0.198 | 0.365 | 0.316 | 0.351 | 0.306 | 0.413 | 0.365 | 0.177 | 0.235 | 0.362 | 0.372 | 0.326 |
| Alborán Sea | 36.00 | -3.96 | 13.21 | 0.033 | 0.100 | 0.067 | 0.000 | 0.033 | 0.036 | 0.000 | 0.036 | 0.192 | 0.033 | 0.033 | 0.000 | 0.033 | 0.000 | 0.150 | 0.000 | 0.033 | 0.000 | 0.033 | 0.133 | 0.033 |
| N. Aegean Sea | 39.76 | 25.02 | 14.74 | 0.000 | 0.085 | 0.085 | 0.047 | 0.000 | 0.009 | 0.010 | 0.028 | 0.290 | 0.000 | 0.010 | 0.000 | 0.000 | 0.000 | 0.265 | 0.000 | 0.000 | 0.000 | 0.000 | 0.183 | 0.000 |
| N. Euboean Gulf | 38.75 | 23.20 | 14.74 | 0.178 | 0.054 | 0.457 | 0.860 | 0.000 | 1.000 | 0.129 | 0.424 | 0.560 | 0.151 | 0.273 | 0.424 | 0.024 | 0.414 | 0.456 | 0.273 | 0.163 | 0.185 | 0.057 | 0.315 | 0.182 |
| Ionian Sea | 38.22 | 22.57 | 14.74 | 0.054 | 0.000 | 0.053 | 0.105 | 0.000 | 0.309 | 0.069 | 0.073 | 0.346 | 0.064 | 0.043 | 0.000 | 0.000 | 0.192 | 0.151 | 0.000 | 0.044 | 0.000 | 0.011 | 0.213 | 0.000 |
| Cretan Sea | 35.65 | 24.99 | 15.79 | 0.000 | 0.034 | 0.032 | 0.081 | 0.000 | 0.011 | 0.000 | 0.000 | 0.109 | 0.000 | 0.012 | 0.000 | 0.000 | 0.000 | 0.185 | 0.000 | 0.000 | 0.000 | 0.000 | 0.170 | 0.012 |

## DISCUSSION

The present study fills the knowledge gap on the population genetic structure of a prospective fisheries target, the Mueller’s pearlside *Maurolicus muelleri*, across most of its distribution range. Three main genetic groups inhabiting the Mediterranean Sea, the Norwegian Fjords and the Atlantic Ocean, respectively supports our hypothesis of three habitat-driven units.

### Genetic markers

The genetic markers used in the current study stem from a two-step procedure: first, mining SNPs from pool sequencing data, and then second: genotyping a subset of selected loci in several hundred individuals using a relatively affordable and high-throughput SNP genotyping method. This same methodological approach was recently used to mine 121 SNPs that revealed remarkable differentiation across habitats in the glacier lanternfish *B. glaciale* (Quintela et al., 2024). In a similar manner, but using ddRAD sequencing, 91 SNPs mined for European sprat unravelled patterns of differentiation (Quintela et al., 2020) later confirmed by full genome sequencing (Petterson et al., 2024). The success of former experiences, the confirmation of the genetic patterns observed with full genome sequencing with the set of 170 selected SNPs together with the outcome of the Genotype Accumulation Curve (Supplement Fig. S2), made us confidently rely on the discriminating capacity of the SNP array used here.

### Patterns of genetic differentiation

The genetic patterns of differentiation of *M. muelleri* align to a great extent with the ones observed in other mesopelagic fish of similar size, *B. glaciale* (Quintela et al., 2024), and add to the body of literature of genetically-identified ecotypic differentiation detected in marine taxa and driven by environmental factors such as bathymetry as in tusk (*Brosme brosme*) (Knutsen et al., 2009) or beaked redfish (*Sebastes mentella*) (Benestan et al., 2021), salinity as in European sprat (*Sprattus sprattus*) (Quintela et al., 2020; Pettersson et al., 2024), or the distinction between marine *vs*. coastal habitat as demonstrated in European anchovy (*Engraulis encrasicolus*) (Le Moan et al., 2016), long-snouted seahorse (*Hyppocampus guttulatus*) (Riquet et al., 2019; Meyer et al., 2024), northern shrimp (*Pandalus borealis*) (Hansen et al., 2021) or Atlantic cod (*Gadus morhua*) (Knutsen et al., 2018).

A major difference between *M. muelleri* and *B. glaciale* is that in the latter, loci in linkage disequilibrium drive a three clusters’ pattern detectable through PCA that matches the genetic signature associated to chromosomal inversions. The arrangement of this putative inversion shows remarkable frequency differences between the Atlantic and the Mediterranean hindering the recombination after gene flow and thus preventing the development of an admixture gradient in that part of the genome. No signature of Mediterranean genetic profile is detected in the genetically uniform Atlantic Ocean samples. However, while we cannot rule out the existence of small chromosome inversions in the genome of *M. muelleri* that could have eluded the scrutiny of our markers, no striations were detected in the PCA and no patterns of differentiation corresponding to different haplotypes were revealed through unsupervised STRUCTURE analyses either. The lack of an apparent inversion in *M. muelleri* is manifested in the Eastern Atlantic samples through an allele frequency cline extending from North Moroccan waters northwards 47 °N. Clinal patterns of differentiation between habitats are also known to the transition Baltic Sea-Atlantic ocean (reviewed in Johannesson & Andre, 2006).

In connection with this putative inversion, the Mediterranean samples of *B. glaciale* displayed less than half the genetic variation compared to the samples from the Atlantic and Norwegian fjords, whereas in *M. muelleri* the Mediterranean samples displayed slightly lower genetic diversity than the rest. This finding highlights the long-term isolation of the Mediterranean unit, which may have led to an adaptation process that reduced their genetic composition to include only alleles suitable to the environment. The values of observed and expected heterozygosity found in this study were similar to *B. glaciale* (Quintela et al., 2024). In contrast, Rodriguez-Ezpeleta et al. (2017) reported exceptionally low levels of genetic diversity (some 4-fold lower) in *M. muelleri* from the Bay of Biscay genotyped at thousands of RAD-seq SNPs, summoning the Wahlund effect. In our study, an extremely low but significant differentiation was detected between the Cantabrian Sea and the Bay of Biscay, while similar patterns were seen in STRUCTURE barplots consistent with the findings of Rodriguez-Ezpeleta et al. (2017). The geographic range of our study allows us to suggest that at K=2, the influence of the Mediterranean genetic profile could account not only for the substructure proposed by Rodriguez-Ezpeleta et al. (2017), but would seem to drive the patterns of differentiation in the Atlantic cluster in a similar manner as the introgression with North East Artic Cod shapes population structure in Norwegian Coastal Cod (Dahle et al., 2018; Jorde et al., 2021). Genetic breaks between the Atlantic and the Mediterranean have been identified in multiple taxa of fish (Bargelloni et al., 2003; Tine et al., 2014; Barros-García et al., 2020; Quintela et al., 2020) including seahorses (Riquet et al., 2019; Meyer et al., 2024), as well as in sponges (Riesgo et al., 2019), molluscs (Pérez-Losada et al., 2002; Lapègue et al., 2023), crustaceans (Reuschel et al., 2010; Jenkins et al., 2019) and harbour porpoise (Fontaine, 2016).

In *M. muelleri* oceanic samples, the Northern Atlantic cluster displays almost no differentiation across the Atlantic from Norway 63 °N to Flemish Cap (3570 km distance in a longitude range from 3.78 °E to 45 °W), which closely aligns with the lack of structure displayed by *B. glaciale* across the Atlantic over distances up to 3940 km (Quintela et al., 2024). Likewise, no genetic structure at mitochondrial cytochrome b was displayed in the North Pacific Ocean in the myctophids *Diaphus theta*, *Stenobrachius leucopsarus*, *S. nannochir* displaying different diel vertical migration patterns (Kojima et al., 2009). Lack of microsatellite genetic differentiation coupled with large genetic diversity was also reported for the southern hemisphere myctophid *Electrona antarctica* (Van de Putte et al., 2012). The homogenizing effect of the Southern Coastal Current in the Southern Ocean not only affects this species, but also others such as humped rockcod *Gobionotothen gibberifrons* (Matschiner et al., 2009) or Antarctic silverfish *Pleuragramma antarctica* (Zane et al., 2006) for which gene flow mediated by larval dispersal resulted in weak or absent genetic structure. In *M. muelleri*, the only northern deviating sample was Vesterålen, which seemed to consist of a physical mixture of genetically distinct fjord and offshore individuals. It could be hypothesized that individuals overcoming the fjord sills could be drifted northwards by the Norwegian Current and retained in Vesterålen area, eventually in combination with the effect of the intensive upwelling that brings deep-see water to Lofoten and Vesterålen (Falk & Nøst, 2013). Otherwise, the differentiation detected in the Atlantic Ocean restricts to the Eastern Southern samples and is driven by the influence of the Mediterranean Sea.

Very little genetic differentiation was observed among samples from the Norwegian fjords, where extremely low but significant differentiation was only detected between the two most distant sites (Osterfjord vs. Boknafjord, *F*_ST_=0.007**). Our result is in agreement with data from an earlier study by Watkins et al. (1996) who reported a lack of genetic structure in *M. muelleri* in the western Norwegian fjords using allozymes. Also with allozymes, Kristoffersen and Salvanes (2009) did not detect differentiation for *B. glaciale* among the fjords, which contrasts with nearly half of the pairwise fjord comparisons revealing significant structure when genotyped with SNPs (Quintela et al., 2024) and with Suneetha and Salvanes (2001) where genetic homogeneity was only detected in fjords with sill depths exceeding 130 m. The sills, typically present at the mouths of the fjords that hinder deep-water exchange with the adjoining coastal areas, could be acting as a physical barrier limiting gene flow within *B. glaciale* (Kristoffersen & Salvanes, 2009), however, they do not seem to have any effect on *M. muelleri*. Differential swimming capacity of both species could be partially responsible for this difference in genetic structure. Tracking of individuals in deep water showed that *B. glaciale* was conspicuously inactive and drifted back and forth with weak tidal currents, essentially acting as plankton while active swimming occasionally occurred in the vertical direction with speeds ranging between <0.5–1 body length s^−1^ (Kaartvedt et al., 2009). In contrast, *M. muelleri* swimming capacity has been reported to reach speeds of 2–7 body length·s^-1^ (*i.e.* 8–30 cm·s^-1^) (Torgersen & Kaartvedt, 2001). In addition to larger swimming capacity, *M. muelleri* has been reported to stay in upper levels in the water column than *B. glaciale* both during day and night time, not only in the Norwegian fjords but also in the Mediterranean and the Atlantic coast (Giske et al., 1990; Staby et al., 2011; Bañón et al., 2016; Rábade Uberos et al., 2021; Olivar et al., 2022; Bernal et al., 2023; Kapelonis et al., 2023) even reaching the surface in the Norwegian fjords during night time (Staby et al., 2011). Furthermore, *M. muelleri* has the capability of changing its behaviour in response to ontogeny and internal state (satiation and hunger) as well as to external stimuli (Staby et al., 2011).

In addition, or maybe because of the reported genetic differentiation across habitats, life history of *M. muelleri* is known to differ geographically, with maximum age reached in Norwegian waters (Gjøsæter & Kawaguchi, 1980). Along the Norwegian coast, the hydrological and topographic characteristics of the fjords may create unique habitat conditions (Farmer & Freeland, 1983) and differentiation between offshore and fjords was detected in traits such as sexual dimorphism (Kristoffersen & Salvanes, 2001), maturation stage, gonadal investment, relative fecundity and growth rate, which displayed increasing trends from the Norwegian Sea to the coast and towards the fjords (Salvanes & Stockley, 1996). The reproductive investment was higher in a fjord with higher predation risk (Bjelland, 1995) and fjord fish also matured earlier in the season than offshore (Salvanes & Stockley, 1996). Mortality rate was reported higher in offshore areas than within the fjords, which was attributed to fish occupying a depth with higher light levels and therefore more vulnerable to predators (Kristoffersen & Salvanes, 1998). Differences in growth, condition and gonad weight indicated different resource levels caused by different population densities (Kristoffersen & Salvanes, 1998). However, back in the day, life history differences were suggested to be phenotypic and not genetically-driven due to the lack of variation fjord-offshore displayed by allozymes (Watkins et al., 1996), which contrasts with the significant differentiation detected with SNPs in this study (average *F*_ST_≈0.09). Interestingly, in the same paper, Watkins et al. (1996) detected fjord-offshore differentiation in *B. glaciale* later confirmed by SNPs (Quintela et al., 2024). Fjord vs. off-shore differentiation has also been described in other marine taxa with high dispersal capacity such as, *e.g.,* tunicate *Ciona intestinalis* (Johannesson et al., 2018), northern shrimp *Pandalus borealis* (Hansen et al., 2021), haddock *Melanogrammus aeglefinus* (Berg et al., 2021), Atlantic cod (Ruzzante et al., 1997; Westgaard & Fevolden, 2007; Pampoulie et al., 2011) or European sprat (Quintela et al., 2020; Pettersson et al., 2024).

Genetic differentiation across areas also mirrors the differentiation found in life history traits in *M. muelleri* from the Bay of Biscay, Celtic Sea, and around 59–63 °N in the Norwegian Sea where length-weight relationships revealed differences associated with the fish’s origin, paralleling the annual and daily otolith growth. Likewise, von Bertalanffy growth models parameters increased progressively northwards, in accordance with Bergmann’s rule (Alvarez et al., 2024). The authors suggested the existence of separated units, either genetically or morphologically, representing differences in biological parameters as a signal of geographical divergence. Our genetic data confirms that the Bay of Biscay significantly differed from the Celtic Sea as well as from Norwegian coastal samples within the same latitudinal range; likewise, the Celtic Sea differed from one of the Norwegian samples.

Oceanographic features like current patterns and discontinuities shape the genetic connectivity in the Mediterranean Sea (Galarza et al., 2009a; Schunter et al., 2011) with the Almería-Oran Front (AOF) being the major point of genetic break between the Atlantic Ocean and the Mediterranean (Patarnello et al., 2007). In addition, the entry of less saline water from the Atlantic through the shallow Strait of Gibraltar represents a barrier to gene flow for multiple species (Galarza et al., 2009a; Galarza et al., 2009b; Marie et al., 2016) and could account for the much lower mesopelagic fish diversity found in the Mediterranean compared with the adjacent Atlantic waters (Olivar et al., 2022). Along the Greek coastline, the Ionian and Aegean Seas shape a complex ecosystem combining a highly irregular coastline and semi-isolated deep basins leading to local differences in species composition (Somarakis et al., 2011; Kapelonis et al., 2023). Genetic differentiation within species has thus been attributed to a combination of historic demographic processes as well as hydrological and ecological traits (see Sarropoulou et al., 2022 and references therein). *M. muelleri* is not oblivious to these factors, which drive the large and highly significant differentiation detected across the Mediterranean (*F_S_*_T_ ranging from 0.05 to 0.18) except for the pair N. Aegean Sea and Cretan Sea, the only samples that do not seem to experience conspicuous physical barriers to gene flow (see Fig. 1). The Gulf of Corinth (Ionian Sea) significantly differed from all the rest (*F_S_*_T_ ranging from 0.05 to 0.06) as it is semi-closed body water with limited communication to the open sea via shallow and narrow channels, which configurates unique hydrological and topographical traits (Drakopoulos & Lascaratos, 1998; Ramfos et al., 2005). The isolation of this site has formerly revealed in both *M. muelleri* and *B. glaciale* by high number of private mtDNA alleles traits (Sarropoulou et al., 2022), in nuclear markers in *B. glaciale* (Quintela et al., 2024), and in the small-spotted catshark S*cyliorhinus canicula* (Kousteni et al., 2015). Among all Mediterranean samples, genetic connectivity seems extremely hampered towards the N Euboean Gulf (*F_S_*_T_ ranging from 0.16 to 0.18), sample that is also genetically closer to the fjord and Atlantic ones.

### Outlier detection and environmental association analyses

The identification of 20 SNPs as candidate outliers to positive selection, alongside 25 candidates to balancing selection highlights the complexity of selective pressures acting on different loci. Adaptive traits relevant to specific environmental contexts are likely maintained through positive selection whereas loci under balancing selection promote genetic diversity, possibly enabling populations to adapt to fluctuating environmental conditions. This distinction emphasizes the necessity of targeting diverse loci when studying adaptation, as it underscores the multifaceted nature of evolutionary pressures in different habitats.

The neutral markers retained the patterns of habitat differentiation observed in the total dataset thus highlighting their role of in shaping genetic structure across habitats. Although PCA and DAPC plots revealed differentiation among habitats, their limitations in capturing the Atlantic-Mediterranean transition indicate a pivotal role of historical factors, such as gene flow and connectivity. This suggests that neutral genetic markers can provide valuable but partial insights into the evolutionary history of the populations. The reduced discrimination capacity signals that more nuanced factors beyond simple neutrality may shape genetic differentiation and warrants further investigation into historical influences.

The candidate loci to positive selection did not have the ability to discriminate between fjords and oceanic samples from the Northern Atlantic cluster. This could be explained by 95% of the candidate outliers being related to temperature according to LFMM and showing extremely similar allele frequencies in the geographic range covering from Vesterålen southwards to the Celtic Sea and westwards to Flemish Cap including both fjord and ocean samples. Likewise, allele frequency clines revealed in 42% of the loci analysed in this study suggest variations due to environmental factors along the latitudinal gradient. In addition to loci linked to temperature, RDA identified a locus seemingly linked to chlorophyll concentration, in agreement with Alamanellis-Zisimopoulos (2023) who reported that bottom depth was the variable with largest importance, followed by the surface chlorophyll concentration (mg/m^3^) and sea current geostrophic velocity (m/s) when modelling the potential habitat of the species in the Greek seas.

An interesting feature is the deviating allele frequencies found in the Greek sample from the N. Euboean Gulf, which tended towards the range of values found in the fjords-Atlantic in contrast with the remaining Mediterranean samples. This pattern occurred in some of the candidate loci to positive selection/linked to temperature while being absent in the neutral ones and emphasizes the potential importance of environmental adaptation. The N. Euboean Gulf is a ∼450 m deep basin that connects with adjacent seas through shallow channels and host only two mesopelagic fish species (*M. muelleri* and *B. glaciale*). The temperature is some ≈2 °C lower than in other Mediterranean regions and it is the only Greek basin that features hypoxic conditions at depths deeper than 350 m (see Suppl. Fig. S12). The lower temperature in combination with the hypoxic conditions and the geographic isolation of the gulf could account for the observed patterns.

Geographic distance explained from 58% (neutral loci) to 61% (all markers) of the genetic differentiation under an Isolation-by-Distance (IBD) pattern for the full set of samples. Likewise, temperature correlated with genetic differentiation driving an Isolation-by-Environment (IBE) pattern both at neutral loci, candidate loci to positive selection and all loci (Suppl. Table S6). However, the correlation between geographic distance and temperature (r_xy_=0.79, *P*<0.001) makes it difficult to disentangle the effect of both variables. Partial Mantel tests correcting IBD by temperature were significant for all loci, whereas when correcting IBE by geographic distance no correlation whatsoever was found at neutral markers suggesting that the relationship between temperature and genetic differentiation disappears when accounting for geographic distance. Candidate outliers to positive selection retained significant when correcting by geographic distance, what highlights the effect of temperature in habitat-driven differentiation in *M. muelleri*.

Genetic response to environmental changes can happen at a high pace in the marine realm. One century was enough to generate a distinct population of European sprat after colonizing an artificially created brackish lake (Quintela et al., 2021; Pettersson et al., 2024). The remarkable response of *M. muelleri* to variable thermal environments suggests that understanding these dynamics may help predict how the species will respond to future climate scenarios and highlights that local adaptation will be critical in informing conservation strategies in the face of ongoing climate change.

### Management implications

Although strongly debated (Pauly et al., 2021), the ambitious estimates of 10 Gt of mesopelagic fish biomass, which is equivalent to ∼100 times the annual catch of all existing fisheries (Hidalgo & Browman, 2019) fuels the interest of harvesting this relatively intact putative resource, mainly as animal feed for aquaculture (Alvheim et al., 2020; Grimaldo et al., 2020). Thus, trial fisheries for mesopelagic fish (including *M. muelleri*) have recently been conducted by Norwegian vessels in the North Atlantic (Standal & Grimaldo, 2021). In the face of an eventual commercial exploitation, and to ensure sustainability, a compelling body of literature highlights the importance of accurately aligning genetic and biological units when defining fisheries stocks (e.g. Allendorf et al., 2008; Hauser & Carvalho, 2008; Waples et al., 2008; Reiss et al., 2009; Funk et al., 2012; Cadrin et al., 2014; Bernatchez et al., 2017; Kerr et al., 2017). The finding of three distinct genetic populations of *M. muelleri* in the North Atlantic and Mediterranean basins provides background information while suggesting that demographic properties should also be mapped to provide a more comprehensive stock outline. Mesopelagic fishes provide multiple ecosystem services, such as regulation ecosystem services, due to their role in the Biological Carbon Pump and carbon sequestration (John et al., 2016), or supporting ecosystem services, since they are prey for a diversity of predators, including economically important species (Iglesias et al., 2023). In consequence, the ecological importance of *M. muelleri* (as of many other mesopelagic fish species) must be thoroughly addressed and put into perspective before attempting any harvest with potential harmful consequences.

## DATA AVAILABILITY

The genotype raw data used in this study can be publicly accessed from the electronic archive of the Institute of Marine Research upon acceptance.

## BENEFIT-SHARING STATEMENT

Benefits generated: A research collaboration was developed with all the scientists from the countries providing genetic samples, all collaborators are included as co-authors, the results of research have been shared with the provider communities and the broader scientific community, and the research addresses a priority concern, in this case the conservation of organisms being studied. More broadly, our group is committed to international scientific partnerships, as well as institutional capacity building.

## ACKNOWLEDGEMENTS

This study was primarily funded by the Norwegian Department for Trade and Fisheries, and further supported by the EU through funding for the MEESO project through the EU H2020 Research and Innovation Programme, Grant Agreement No 817669. Samples from the Western Mediterranean were collected during a survey performed within the SUMMER EU H2020 Research and Innovation Programme, Grant Agreement No 817806. Samples from the Eastern Mediterranean were obtained under the project MesoBED, funded by the Hellenic Foundation for Research and Innovation and the General Secretariat of Research and Innovation (Greece) (Project No. 449). The sample from Moroccan waters was collected through the scientific surveys with the research vessel *Dr. Fridtjof Nansen* as part of the collaboration between the Food and Agriculture Organization of the United Nations (FAO) on behalf of the EAF-Nansen Programme and the Kingdom of Morocco. The EAF-Nansen Programme is a partnership between the FAO, the Norwegian Agency for Development Cooperation (Norad), and the Institute of Marine Research (IMR) in Norway for sustainable management of the fisheries in partner countries and regions. The sample from the Flemish Cap was collected by CISC (Spain) on behalf of the European Union sampling programme during a survey conducted onboard the research vessel *Vizconde de Eza*. We are grateful to Nikolaos Nikolioudakis (IMR) and Iñaki Mendibil (AZTI) for their valuable help with sampling logistics.

