## Supplement for "Genetics in the ocean’s twilight zone: Population structure of the Mueller’s pearlside across its distribution range"

**TABLES**

**Table S1.-** List of the 30 discarded SNPs together with the reasons for them to be removed.

| **Locus** | **Reasons to remove** |
| --- | --- |
| P325 | 100% missing loci overall |
| P378 | 100% missing loci overall |
| P437 | 100% missing loci overall |
| P537 | 100% missing loci overall |
| P614 | 100% missing loci overall |
| P688 | 100% missing loci overall |
| P834 | 100% missing loci overall |
| P1153 | 100% missing loci overall |
| P1284 | 100% missing loci overall |
| P1295 | 100% missing loci overall |
| P1423 | 100% missing loci overall |
| P1479 | 100% missing loci overall |
| P1638 | 100% missing loci overall |
| P1675 | 100% missing loci overall |
| P1774 | 100% missing loci overall |
| P1775 | 100% missing loci overall |
| P1869 | 100% missing loci overall |
| P1900 | 100% missing loci overall |
| P1953 | 100% missing loci overall |
| P660 | 66.3% missing loci overall |
| P974 | 70.2% missing loci overall |
| P1784 | 81.35% missing loci overall |
| P1246 | Empty in various samples |
| P1251 | Empty in various samples |
| P1315 | Empty in various samples |
| P1346 | Empty in various samples |
| P1520 | Empty in various samples |
| P1568 | Empty in various samples |
| P1832 | Empty in various samples |
| P1922 | Empty in various samples |

**Table S2.-** Genetic differentiation between geographically explicit samples assessed with 170 SNPs: Heatmap of pairwise *F*_ST_ values in the bottom diagonal and corresponding *P*-values after 10,000 permutations in the top diagonal, with the ones significantly different from zero after FDR correction highlighted in boldface type. Greener colours indicate low differentiation increasing towards red to indicate larger differentiation. Sample names shaded in grey have sampling sizes ranging from 10 to 18 individuals.

|  | **Osterfj** | **Byfj** | **Korsfj** | **Bjørnafj** | **Boknafj** | **Vesterål** | **Norway63N** | **Iceland** | **Norway60N** | **Norway59N** | **AtlanticXIIc** | **Celtic** | **Flemish** | **Atlantic47N** | **Biscay Gulf** | **Cantabrian** | **Atlantic40N** | **Morocco** | **Alborán** | **NAegean** | **NEuboean** | **Ionian** | **Cretan** |
| --- | --- | --- | --- | --- | --- | --- | --- | --- | --- | --- | --- | --- | --- | --- | --- | --- | --- | --- | --- | --- | --- | --- | --- |
| **Osterfj** | * | 0.855 | 0.830 | 0.752 | **0.002** | **0.030** | **0.000** | **0.000** | **0.000** | **0.000** | **0.000** | **0.000** | **0.000** | **0.000** | **0.000** | **0.000** | **0.000** | **0.000** | **0.000** | **0.000** | **0.000** | **0.000** | **0.000** |
| **Byfj** | 0.000 | * | 0.882 | 0.979 | 0.981 | 0.141 | **0.000** | **0.000** | **0.000** | **0.000** | **0.000** | **0.000** | **0.000** | **0.000** | **0.000** | **0.000** | **0.000** | **0.000** | **0.000** | **0.000** | **0.000** | **0.000** | **0.000** |
| **Korsfj** | 0.000 | 0.000 | * | 0.879 | 0.491 | 0.139 | **0.000** | **0.000** | **0.000** | **0.000** | **0.000** | **0.000** | **0.000** | **0.000** | **0.000** | **0.000** | **0.000** | **0.000** | **0.000** | **0.000** | **0.000** | **0.000** | **0.000** |
| **Bjørnafj** | 0.000 | 0.000 | 0.000 | * | 0.571 | **0.013** | **0.000** | **0.000** | **0.000** | **0.000** | **0.000** | **0.000** | **0.000** | **0.000** | **0.000** | **0.000** | **0.000** | **0.000** | **0.000** | **0.000** | **0.000** | **0.000** | **0.000** |
| **Boknafj** | 0.007 | 0.000 | 0.000 | 0.000 | * | **0.004** | **0.000** | **0.000** | **0.000** | **0.000** | **0.000** | **0.000** | **0.000** | **0.000** | **0.000** | **0.000** | **0.000** | **0.000** | **0.000** | **0.000** | **0.000** | **0.000** | **0.000** |
| **Vesterål** | 0.009 | 0.005 | 0.005 | 0.010 | 0.012 | * | **0.000** | **0.000** | **0.000** | **0.000** | **0.000** | **0.000** | **0.001** | **0.000** | **0.000** | **0.000** | **0.000** | **0.000** | **0.000** | **0.000** | **0.000** | **0.000** | **0.000** |
| **Norway63N** | 0.086 | 0.096 | 0.081 | 0.081 | 0.096 | 0.048 | * | 0.162 | 0.999 | 0.992 | 0.721 | 0.663 | 0.050 | **0.000** | **0.000** | **0.000** | **0.000** | **0.000** | **0.000** | **0.000** | **0.000** | **0.000** | **0.000** |
| **Iceland** | 0.104 | 0.117 | 0.102 | 0.102 | 0.121 | 0.065 | 0.012 | * | 0.519 | 0.175 | 0.552 | 0.402 | 0.594 | **0.000** | **0.000** | **0.000** | **0.000** | **0.000** | **0.000** | **0.000** | **0.000** | **0.000** | **0.000** |
| **Norway60N** | 0.089 | 0.091 | 0.082 | 0.088 | 0.102 | 0.054 | 0.000 | 0.000 | * | 1.000 | 0.821 | **0.026** | 0.345 | **0.000** | **0.000** | **0.000** | **0.000** | **0.000** | **0.000** | **0.000** | **0.000** | **0.000** | **0.000** |
| **Norway59N** | 0.083 | 0.086 | 0.077 | 0.082 | 0.096 | 0.042 | 0.000 | 0.005 | 0.000 | * | 0.997 | 0.959 | **0.008** | **0.000** | **0.000** | **0.000** | **0.000** | **0.000** | **0.000** | **0.000** | **0.000** | **0.000** | **0.000** |
| **AtlanticXIIc** | 0.094 | 0.096 | 0.089 | 0.094 | 0.108 | 0.054 | 0.000 | 0.000 | 0.000 | 0.000 | * | 0.251 | 0.075 | **0.000** | **0.000** | **0.000** | **0.000** | **0.000** | **0.000** | **0.000** | **0.000** | **0.000** | **0.000** |
| **Celtic** | 0.079 | 0.082 | 0.075 | 0.080 | 0.093 | 0.046 | 0.000 | 0.001 | 0.004 | 0.000 | 0.001 | * | 0.316 | **0.000** | **0.000** | **0.000** | **0.000** | **0.000** | **0.000** | **0.000** | **0.000** | **0.000** | **0.000** |
| **Flemish** | 0.087 | 0.090 | 0.082 | 0.080 | 0.103 | 0.049 | 0.019 | 0.000 | 0.002 | 0.013 | 0.007 | 0.002 | * | **0.000** | **0.000** | **0.000** | **0.000** | **0.000** | **0.000** | **0.000** | **0.000** | **0.000** | **0.000** |
| **Atlantic47N** | 0.109 | 0.106 | 0.103 | 0.106 | 0.114 | 0.074 | 0.041 | 0.052 | 0.046 | 0.045 | 0.049 | 0.031 | 0.055 | * | 0.387 | **0.000** | 0.454 | **0.000** | **0.000** | **0.000** | **0.000** | **0.000** | **0.000** |
| **Biscay Gulf** | 0.122 | 0.112 | 0.117 | 0.119 | 0.122 | 0.088 | 0.066 | 0.078 | 0.078 | 0.080 | 0.082 | 0.056 | 0.090 | 0.000 | * | **0.002** | 0.996 | **0.000** | **0.000** | **0.000** | **0.000** | **0.000** | **0.000** |
| **Cantabrian** | 0.176 | 0.172 | 0.169 | 0.169 | 0.176 | 0.147 | 0.123 | 0.130 | 0.129 | 0.131 | 0.132 | 0.104 | 0.139 | 0.025 | 0.011 | * | **0.005** | **0.021** | **0.000** | **0.000** | **0.000** | **0.000** | **0.000** |
| **Atlantic40N** | 0.122 | 0.115 | 0.116 | 0.120 | 0.126 | 0.093 | 0.067 | 0.075 | 0.076 | 0.072 | 0.080 | 0.056 | 0.080 | 0.000 | 0.000 | 0.009 | * | **0.000** | **0.000** | **0.000** | **0.000** | **0.000** | **0.000** |
| **Morocco** | 0.213 | 0.206 | 0.207 | 0.209 | 0.214 | 0.185 | 0.174 | 0.183 | 0.178 | 0.178 | 0.182 | 0.150 | 0.184 | 0.051 | 0.029 | 0.005 | 0.025 | * | **0.000** | **0.000** | **0.000** | **0.000** | **0.000** |
| **Alborán** | 0.338 | 0.348 | 0.334 | 0.331 | 0.342 | 0.339 | 0.363 | 0.370 | 0.322 | 0.334 | 0.327 | 0.286 | 0.372 | 0.162 | 0.128 | 0.080 | 0.115 | 0.043 | * | **0.000** | **0.000** | **0.000** | **0.000** |
| **NAegean** | 0.360 | 0.357 | 0.352 | 0.347 | 0.354 | 0.355 | 0.370 | 0.391 | 0.349 | 0.358 | 0.355 | 0.326 | 0.390 | 0.212 | 0.177 | 0.125 | 0.168 | 0.091 | 0.062 | * | **0.000** | **0.000** | 0.516 |
| **NEuboean** | 0.214 | 0.214 | 0.206 | 0.208 | 0.214 | 0.191 | 0.198 | 0.213 | 0.194 | 0.195 | 0.199 | 0.177 | 0.209 | 0.108 | 0.093 | 0.072 | 0.087 | 0.084 | 0.160 | 0.174 | * | **0.000** | **0.000** |
| **Ionian** | 0.327 | 0.323 | 0.321 | 0.318 | 0.323 | 0.319 | 0.337 | 0.355 | 0.321 | 0.330 | 0.326 | 0.295 | 0.353 | 0.181 | 0.146 | 0.110 | 0.144 | 0.074 | 0.063 | 0.049 | 0.133 | * | **0.000** |
| **Cretan** | 0.365 | 0.364 | 0.357 | 0.351 | 0.362 | 0.367 | 0.382 | 0.402 | 0.352 | 0.360 | 0.359 | 0.330 | 0.401 | 0.215 | 0.185 | 0.129 | 0.171 | 0.097 | 0.075 | 0.000 | 0.179 | 0.062 | * |

**Table S3.-** Genetic differentiation between geographically explicit samples assessed with 78 putatively neutral loci: Heatmap of pairwise *F*_ST_ values in the bottom diagonal and corresponding *P*-values after 10,000 permutations in the top diagonal, with the ones significantly different from zero after FDR correction highlighted in boldface type. Greener colours indicate low differentiation increasing towards red to indicate larger differentiation. Sample names shaded in grey have sampling sizes ranging from 10 to 18 individuals.

|  | **Osterfj** | **Byfj** | **Korsfj** | **Bjørnafj** | **Boknafj** | **Vesterål** | **Norway63N** | **Iceland** | **Norway60N** | **Norway59N** | **AtlanticXIIc** | **Celtic** | **Flemish** | **Atlantic47N** | **Biscay Gulf** | **Cantabrian** | **Atlantic40N** | **Morocco** | **Alborán** | **NAegean** | **NEuboean** | **Ionian** | **Cretan** |
| --- | --- | --- | --- | --- | --- | --- | --- | --- | --- | --- | --- | --- | --- | --- | --- | --- | --- | --- | --- | --- | --- | --- | --- |
| **Osterfj** | * | 0.960 | 0.307 | 0.859 | **0.016** | **0.005** | **0.000** | **0.000** | **0.000** | **0.000** | **0.000** | **0.000** | **0.000** | **0.000** | **0.000** | **0.000** | **0.000** | **0.000** | **0.000** | **0.000** | **0.000** | **0.000** | **0.000** |
| **Byfj** | 0.000 | * | 1.000 | 1.000 | 1.000 | 0.354 | **0.000** | **0.000** | **0.000** | **0.000** | **0.000** | **0.000** | **0.000** | **0.000** | **0.000** | **0.000** | **0.000** | **0.000** | **0.000** | **0.000** | **0.000** | **0.000** | **0.000** |
| **Korsfj** | 0.001 | 0.000 | * | 0.597 | 0.672 | 0.069 | **0.000** | **0.000** | **0.000** | **0.000** | **0.000** | **0.000** | **0.000** | **0.000** | **0.000** | **0.000** | **0.000** | **0.000** | **0.000** | **0.000** | **0.000** | **0.000** | **0.000** |
| **Bjørnafj** | 0.000 | 0.000 | 0.000 | * | 0.861 | 0.033 | **0.000** | **0.000** | **0.000** | **0.000** | **0.000** | **0.000** | **0.000** | **0.000** | **0.000** | **0.000** | **0.000** | **0.000** | **0.000** | **0.000** | **0.000** | **0.000** | **0.000** |
| **Boknafj** | 0.006 | 0.000 | 0.000 | 0.000 | * | **0.007** | **0.000** | **0.000** | **0.000** | **0.000** | **0.000** | **0.000** | **0.000** | **0.000** | **0.000** | **0.000** | **0.000** | **0.000** | **0.000** | **0.000** | **0.000** | **0.000** | **0.000** |
| **Vesterål** | 0.017 | 0.003 | 0.009 | 0.012 | 0.016 | * | **0.000** | **0.000** | **0.000** | **0.000** | **0.000** | **0.000** | **0.023** | **0.000** | **0.000** | **0.000** | **0.000** | **0.000** | **0.000** | **0.000** | **0.000** | **0.000** | **0.000** |
| **Norway63N** | 0.100 | 0.117 | 0.092 | 0.101 | 0.109 | 0.059 | * | **0.022** | 0.955 | 0.962 | 0.282 | 0.961 | 0.047 | 0.176 | 0.033 | **0.000** | 0.159 | **0.000** | **0.000** | **0.000** | **0.000** | **0.000** | **0.000** |
| **Iceland** | 0.111 | 0.124 | 0.110 | 0.114 | 0.125 | 0.070 | 0.031 | * | 0.167 | 0.102 | 0.374 | 0.683 | 0.363 | **0.000** | **0.000** | **0.000** | **0.000** | **0.000** | **0.000** | **0.000** | **0.000** | **0.000** | **0.000** |
| **Norway60N** | 0.104 | 0.105 | 0.094 | 0.107 | 0.117 | 0.063 | 0.000 | 0.008 | * | 1.000 | 0.853 | 0.805 | 0.245 | **0.000** | **0.000** | **0.000** | **0.000** | **0.000** | **0.000** | **0.000** | **0.000** | **0.000** | **0.000** |
| **Norway59N** | 0.098 | 0.103 | 0.089 | 0.102 | 0.112 | 0.054 | 0.000 | 0.010 | 0.000 | * | 1.000 | 1.000 | **0.015** | **0.000** | **0.000** | **0.000** | **0.000** | **0.000** | **0.000** | **0.000** | **0.000** | **0.000** | **0.000** |
| **AtlanticXIIc** | 0.117 | 0.119 | 0.106 | 0.118 | 0.129 | 0.069 | 0.002 | 0.001 | 0.000 | 0.000 | * | 0.560 | 0.125 | **0.000** | **0.000** | **0.000** | **0.000** | **0.000** | **0.000** | **0.000** | **0.000** | **0.000** | **0.000** |
| **Celtic** | 0.094 | 0.098 | 0.086 | 0.097 | 0.104 | 0.053 | 0.000 | 0.000 | 0.000 | 0.000 | 0.000 | * | 0.263 | **0.001** | **0.000** | **0.000** | **0.000** | **0.000** | **0.000** | **0.000** | **0.000** | **0.000** | **0.000** |
| **Flemish** | 0.086 | 0.085 | 0.080 | 0.088 | 0.100 | 0.034 | 0.027 | 0.007 | 0.005 | 0.018 | 0.008 | 0.004 | * | **0.001** | **0.000** | **0.000** | **0.002** | **0.000** | **0.000** | **0.000** | **0.000** | **0.000** | **0.000** |
| **Atlantic47N** | 0.101 | 0.110 | 0.093 | 0.101 | 0.104 | 0.060 | 0.005 | 0.042 | 0.017 | 0.014 | 0.021 | 0.012 | 0.037 | * | 0.127 | **0.001** | 0.890 | **0.000** | **0.000** | **0.000** | **0.000** | **0.000** | **0.000** |
| **Biscay Gulf** | 0.100 | 0.099 | 0.092 | 0.098 | 0.096 | 0.056 | 0.014 | 0.043 | 0.038 | 0.042 | 0.046 | 0.024 | 0.056 | 0.005 | * | 0.324 | 0.973 | **0.015** | **0.000** | **0.000** | **0.000** | **0.000** | **0.000** |
| **Cantabrian** | 0.118 | 0.122 | 0.110 | 0.112 | 0.113 | 0.085 | 0.039 | 0.070 | 0.057 | 0.064 | 0.061 | 0.047 | 0.075 | 0.015 | 0.002 | * | 0.093 | 0.450 | **0.000** | **0.000** | **0.000** | **0.000** | **0.000** |
| **Atlantic40N** | 0.093 | 0.092 | 0.084 | 0.092 | 0.093 | 0.054 | 0.006 | 0.034 | 0.027 | 0.022 | 0.033 | 0.019 | 0.031 | 0.000 | 0.000 | 0.005 | * | **0.001** | **0.000** | **0.000** | **0.000** | **0.000** | **0.000** |
| **Morocco** | 0.134 | 0.134 | 0.127 | 0.129 | 0.127 | 0.099 | 0.067 | 0.108 | 0.085 | 0.088 | 0.091 | 0.074 | 0.100 | 0.026 | 0.009 | 0.000 | 0.014 | * | **0.000** | **0.000** | **0.000** | **0.000** | **0.000** |
| **Alborán** | 0.161 | 0.182 | 0.161 | 0.157 | 0.157 | 0.154 | 0.193 | 0.236 | 0.174 | 0.185 | 0.173 | 0.155 | 0.221 | 0.109 | 0.075 | 0.054 | 0.074 | 0.035 | * | **0.000** | **0.000** | **0.000** | **0.000** |
| **NAegean** | 0.252 | 0.262 | 0.246 | 0.234 | 0.243 | 0.248 | 0.277 | 0.338 | 0.257 | 0.278 | 0.267 | 0.261 | 0.318 | 0.210 | 0.177 | 0.146 | 0.173 | 0.125 | 0.069 | * | **0.000** | **0.000** | 0.331 |
| **NEuboean** | 0.140 | 0.161 | 0.138 | 0.130 | 0.147 | 0.112 | 0.131 | 0.169 | 0.130 | 0.138 | 0.131 | 0.127 | 0.138 | 0.119 | 0.104 | 0.092 | 0.095 | 0.101 | 0.107 | 0.150 | * | **0.000** | **0.000** |
| **Ionian** | 0.209 | 0.221 | 0.201 | 0.193 | 0.199 | 0.190 | 0.226 | 0.284 | 0.226 | 0.241 | 0.234 | 0.219 | 0.268 | 0.176 | 0.139 | 0.127 | 0.141 | 0.103 | 0.053 | 0.035 | 0.115 | * | **0.000** |
| **Cretan** | 0.252 | 0.266 | 0.247 | 0.235 | 0.246 | 0.256 | 0.285 | 0.342 | 0.256 | 0.274 | 0.266 | 0.258 | 0.326 | 0.210 | 0.180 | 0.149 | 0.172 | 0.130 | 0.081 | 0.001 | 0.158 | 0.054 | * |

**Table S4.-** Genetic differentiation between geographically explicit samples assessed with 20 candidate loci to directional selection: Heatmap of pairwise *F*_ST_ values in the bottom diagonal and corresponding *P*-values after 10,000 permutations in the top diagonal, with the ones significantly different from zero after FDR correction highlighted in boldface type. Greener colours indicate low differentiation increasing towards red to indicate larger differentiation. Sample names shaded in grey have sampling sizes ranging from 10 to 18 individuals.

|  | **Osterfj** | **Byfj** | **Korsfj** | **Bjørnafj** | **Boknafj** | **Vesterål** | **Norway63N** | **Iceland** | **Norway60N** | **Norway59N** | **AtlanticXIIc** | **Celtic** | **Flemish** | **Atlantic47N** | **Biscay Gulf** | **Cantabrian** | **Atlantic40N** | **Morocco** | **Alborán** | **NAegean** | **NEuboean** | **Ionian** | **Cretan** |
| --- | --- | --- | --- | --- | --- | --- | --- | --- | --- | --- | --- | --- | --- | --- | --- | --- | --- | --- | --- | --- | --- | --- | --- |
| **Osterfj** | * | 0.130 | 0.701 | 0.296 | 0.167 | 0.951 | 0.324 | 0.386 | **0.002** | 0.065 | **0.021** | **0.000** | **0.001** | **0.000** | **0.000** | **0.000** | **0.000** | **0.000** | **0.000** | **0.000** | **0.000** | **0.000** | **0.000** |
| **Byfj** | 0.009 | * | 0.389 | 0.354 | 0.095 | 0.430 | 0.489 | 0.187 | **0.025** | 0.822 | 0.309 | **0.000** | **0.001** | **0.000** | **0.000** | **0.000** | **0.000** | **0.000** | **0.000** | **0.000** | **0.000** | **0.000** | **0.000** |
| **Korsfj** | 0.000 | 0.000 | * | 0.929 | 0.635 | 0.998 | 0.919 | 0.723 | 0.070 | 0.925 | 0.374 | **0.000** | **0.020** | **0.000** | **0.000** | **0.000** | **0.000** | **0.000** | **0.000** | **0.000** | **0.000** | **0.000** | **0.000** |
| **Bjørnafj** | 0.004 | 0.002 | 0.000 | * | 0.996 | 0.949 | 0.579 | 0.534 | 0.033 | 0.968 | 0.425 | **0.001** | 0.043 | **0.000** | **0.000** | **0.000** | **0.000** | **0.000** | **0.000** | **0.000** | **0.000** | **0.000** | **0.000** |
| **Boknafj** | 0.008 | 0.013 | 0.000 | 0.000 | * | 0.831 | 0.446 | 0.567 | **0.028** | 0.927 | 0.222 | **0.001** | 0.043 | **0.000** | **0.000** | **0.000** | **0.000** | **0.000** | **0.000** | **0.000** | **0.000** | **0.000** | **0.000** |
| **Vesterål** | 0.000 | 0.000 | 0.000 | 0.000 | 0.000 | * | 0.687 | 0.649 | 0.155 | 0.649 | 0.359 | 0.048 | **0.024** | **0.000** | **0.000** | **0.000** | **0.000** | **0.000** | **0.000** | **0.000** | **0.000** | **0.000** | **0.000** |
| **Norway63N** | 0.002 | 0.000 | 0.000 | 0.000 | 0.000 | 0.000 | * | 0.871 | 0.969 | 0.976 | 0.981 | 0.221 | 0.046 | **0.000** | **0.000** | **0.000** | **0.000** | **0.000** | **0.000** | **0.000** | **0.000** | **0.000** | **0.000** |
| **Iceland** | 0.001 | 0.014 | 0.000 | 0.000 | 0.000 | 0.000 | 0.000 | * | 0.680 | 0.711 | 0.561 | 0.764 | 0.290 | **0.000** | **0.000** | **0.000** | **0.000** | **0.000** | **0.000** | **0.000** | **0.000** | **0.000** | **0.000** |
| **Norway60N** | 0.042 | 0.024 | 0.015 | 0.022 | 0.023 | 0.016 | 0.000 | 0.000 | * | 1.000 | 0.484 | **0.028** | 0.107 | **0.000** | **0.000** | **0.000** | **0.000** | **0.000** | **0.000** | **0.000** | **0.000** | **0.000** | **0.000** |
| **Norway59N** | 0.018 | 0.000 | 0.000 | 0.000 | 0.000 | 0.000 | 0.000 | 0.000 | 0.000 | * | 1.000 | 0.241 | 0.227 | **0.000** | **0.000** | **0.000** | **0.000** | **0.000** | **0.000** | **0.000** | **0.000** | **0.000** | **0.000** |
| **AtlanticXIIc** | 0.025 | 0.002 | 0.001 | 0.001 | 0.006 | 0.003 | 0.000 | 0.000 | 0.000 | 0.000 | * | 0.033 | 0.095 | **0.000** | **0.000** | **0.000** | **0.000** | **0.000** | **0.000** | **0.000** | **0.000** | **0.000** | **0.000** |
| **Celtic** | 0.064 | 0.047 | 0.040 | 0.041 | 0.038 | 0.029 | 0.008 | 0.000 | 0.019 | 0.005 | 0.020 | * | 0.474 | **0.000** | **0.000** | **0.000** | **0.000** | **0.000** | **0.000** | **0.000** | **0.000** | **0.000** | **0.000** |
| **Flemish** | 0.119 | 0.124 | 0.070 | 0.063 | 0.055 | 0.076 | 0.059 | 0.010 | 0.032 | 0.017 | 0.038 | 0.000 | * | **0.000** | **0.000** | **0.000** | **0.000** | **0.000** | **0.000** | **0.000** | **0.000** | **0.000** | **0.000** |
| **Atlantic47N** | 0.313 | 0.276 | 0.292 | 0.304 | 0.305 | 0.236 | 0.215 | 0.167 | 0.256 | 0.258 | 0.259 | 0.154 | 0.193 | * | 0.798 | **0.000** | 0.371 | **0.000** | **0.000** | **0.000** | **0.000** | **0.000** | **0.000** |
| **Biscay Gulf** | 0.399 | 0.343 | 0.380 | 0.395 | 0.400 | 0.301 | 0.289 | 0.229 | 0.350 | 0.357 | 0.348 | 0.232 | 0.269 | 0.000 | * | **0.001** | 0.783 | **0.000** | **0.000** | **0.000** | **0.000** | **0.000** | **0.000** |
| **Cantabrian** | 0.570 | 0.534 | 0.554 | 0.568 | 0.570 | 0.473 | 0.464 | 0.400 | 0.533 | 0.538 | 0.531 | 0.413 | 0.431 | 0.100 | 0.062 | * | **0.001** | **0.008** | **0.000** | **0.000** | **0.000** | **0.000** | **0.000** |
| **Atlantic40N** | 0.377 | 0.332 | 0.360 | 0.372 | 0.374 | 0.301 | 0.285 | 0.238 | 0.334 | 0.335 | 0.334 | 0.236 | 0.269 | 0.001 | 0.000 | 0.043 | * | **0.000** | **0.000** | **0.000** | **0.000** | **0.000** | **0.000** |
| **Morocco** | 0.628 | 0.597 | 0.616 | 0.627 | 0.629 | 0.555 | 0.547 | 0.499 | 0.599 | 0.603 | 0.599 | 0.503 | 0.527 | 0.191 | 0.134 | 0.028 | 0.109 | * | **0.000** | **0.000** | **0.000** | **0.000** | **0.000** |
| **Alborán** | 0.906 | 0.901 | 0.899 | 0.900 | 0.910 | 0.902 | 0.898 | 0.863 | 0.878 | 0.896 | 0.883 | 0.792 | 0.902 | 0.480 | 0.428 | 0.275 | 0.377 | 0.154 | * | 0.435 | **0.000** | **0.004** | 0.813 |
| **NAegean** | 0.889 | 0.882 | 0.884 | 0.885 | 0.891 | 0.879 | 0.875 | 0.857 | 0.869 | 0.880 | 0.872 | 0.812 | 0.874 | 0.561 | 0.521 | 0.349 | 0.454 | 0.204 | 0.000 | * | **0.000** | **0.000** | 0.201 |
| **NEuboean** | 0.619 | 0.582 | 0.605 | 0.616 | 0.619 | 0.555 | 0.537 | 0.489 | 0.584 | 0.589 | 0.588 | 0.487 | 0.525 | 0.182 | 0.157 | 0.070 | 0.119 | 0.138 | 0.439 | 0.495 | * | **0.000** | **0.000** |
| **Ionian** | 0.839 | 0.826 | 0.832 | 0.836 | 0.841 | 0.812 | 0.806 | 0.778 | 0.816 | 0.826 | 0.818 | 0.745 | 0.801 | 0.456 | 0.405 | 0.224 | 0.348 | 0.101 | 0.058 | 0.065 | 0.350 | * | **0.000** |
| **Cretan** | 0.899 | 0.893 | 0.894 | 0.895 | 0.901 | 0.893 | 0.888 | 0.870 | 0.878 | 0.890 | 0.881 | 0.819 | 0.889 | 0.562 | 0.524 | 0.352 | 0.457 | 0.214 | 0.000 | 0.006 | 0.498 | 0.071 | * |

**Table S5.-** Summary of information provided by different approaches for all loci involved in allele frequency clines. Outlier detection: positive selection, balancing selection or no departure from neutrality after consensus between BayeScan and Arlequin. Blue depicts the loci identified as associated to temperature at 200 m depth according to LFMM. Loci associated to environmental factors according to RDA are indicated. Loadings on the first axis of the DAPC highlighted in boldface font corresponds to the loci that contributed the most to the genetic differentiation. Model that fits best each cline according to HZAR. Cline centre per locus with those shaded in grey corresponding to the loci that fit within the limits of the reference cline built with STRUCTURE-Q score. Gene predictions identified using BLAST on the SNP flanking regions.

| **Locus** | **Outlier detection** | **LFMM** | **RDA** | **DAPC_Loadings1axis** | **Cline model** | **Cline centre (km)** | **BLAST** |
| --- | --- | --- | --- | --- | --- | --- | --- |
| P1446 | Positive | Outlier |  | **0.23** | fixR | 107 |  |
| P1691 | Positive | Outlier |  | **0.21** | typN | 2340 | PREDICTED: *Calypte anna* ADAM metallopeptidase domain 12 (ADAM12), transcript variant X1 |
| P635 | Positive | Outlier |  | 0.18 | typL | 2841 |  |
| P1636 | Positive | Outlier |  | **0.27** | optR | 3035 |  |
| P1770 | Positive | Outlier |  | **0.32** | fixR | 3064 |  |
| P1945 | Positive | Outlier |  | **0.26** | fixR | 3157 |  |
| P709 | Positive | Outlier |  | 0.14 | typR | 3160 |  |
| P105 | Positive | Outlier |  | **0.24** | optB | 3162 | *Paramormyrops kingsleyae* disintegrin and metalloproteinase domain-containing protein 12-like |
| P1267 | Positive | Outlier |  | **0.24** | optB | 3176 |  |
| P1569 | Positive | Outlier |  | 0.14 | typR | 3183 |  |
| P942 | Positive | Outlier | Temperature | 0.19 | optM | 3230 | PREDICTED: *Sparus aurata* zinc finger and BTB domain containing 39 (zbtb39) |
| P1458 | Positive | Outlier | Temperature | **0.23** | typR | 3251 |  |
| P676 | Positive | Outlier |  | **0.33** | optN | 3482 |  |
| P1989 | Positive | Outlier |  | **0.23** | optN | 3573 | PREDICTED: *Hypomesus transpacificus* zinc finger protein 609-like (LOC124481548), transcript variant X2 |
| P402 | Positive | Outlier |  | **0.32** | typR | 3588 |  |
| P1500 | Positive | Outlier |  | 0.20 | typN | 3597 |  |
| P1856 | Positive | Outlier |  | **0.23** | typN | 3635 |  |
| P1606 | Positive | Outlier |  | 0.09 | fixR | 3815 |  |
| P989 | Positive | Outlier |  | 0.17 | typR | 4191 | PREDICTED: *Seriola dumerili* MAX interactor 1, dimerization protein (mxi1), transcript variant X2 |
| P1668 | Positive |  |  | 0.06 | optN | 3155 |  |
| P253 | Positive BayeScan | Outlier |  | 0.10 | fixN | 3338 | PREDICTED: *Salvelinus fontinalis* anoctamin 5a (ano5a), transcript variant X2 |
| P359 | Positive BayeScan | Outlier |  | 0.18 | fixR | 3468 |  |
| P318 | Neutral |  |  | 0.12 | fixN | 41 |  |
| P1337 | Neutral |  |  | 0.01 | typM | 396 |  |
| P481 | Neutral |  |  | 0.14 | fixN | 1617 |  |
| P790 | Neutral |  |  | 0.20 | fixN | 1701 |  |
| P1345 | Neutral |  |  | 0.05 | fixN | 2348 |  |
| P1661 | Neutral |  |  | **0.24** | optN | 2634 |  |
| P1133 | Neutral |  |  | 0.13 | optN | 2670 |  |
| P1010 | Neutral |  |  | 0.05 | fixN | 2732 |  |
| P449 | Neutral |  |  | 0.03 | fixN | 2743 |  |
| P267 | Neutral |  |  | 0.02 | optN | 2778 |  |
| P045 | Neutral |  |  | 0.16 | fixR | 2778 |  |
| P1863 | Neutral |  |  | **0.21** | fixR | 2782 |  |
| P102 | Neutral |  |  | 0.02 | optN | 3036 |  |
| P296 | Neutral |  |  | 0.06 | optN | 3053 |  |
| P1883 | Neutral |  |  | 0.03 | fixN | 3144 |  |
| P1682 | Neutral |  |  | 0.01 | fixN | 3170 |  |
| P1833 | Neutral |  |  | 0.11 | fixN | 3342 |  |
| P1241 | Neutral |  |  | 0.07 | fixN | 3376 |  |
| P661 | Neutral |  |  | 0.02 | fixN | 3395 |  |
| P303 | Neutral |  |  | 0.20 | fixN | 3403 |  |
| P356 | Neutral |  |  | 0.05 | fixN | 3417 |  |
| P1930 | Neutral |  |  | 0.00 | fixN | 3452 |  |
| P307 | Neutral |  |  | 0.20 | optN | 3477 |  |
| P793 | Neutral |  |  | 0.04 | fixN | 3625 |  |
| P1981 | Neutral |  |  | 0.13 | fixN | 3631 |  |
| P1757 | Neutral |  |  | 0.05 | fixN | 3654 |  |
| P643 | Neutral |  |  | 0.08 | fixN | 3686 |  |
| P479 | Neutral |  |  | 0.04 | fixN | 3762 |  |
| P545 | Neutral |  |  | 0.03 | fixN | 3784 |  |
| P488 | Neutral |  |  | 0.11 | fixN | 3971 |  |
| P002 | Neutral |  |  | 0.16 | optN | 4291 |  |
| P584 | Neutral |  |  | 0.05 | optN | 4302 |  |
| P1062 | Neutral |  |  | **0.25** | optN | 4546 |  |
| P654 | Neutral |  |  | 0.04 | fixN | 4656 |  |
| P1768 | Neutral |  |  | **0.22** | fixN | 4682 |  |
| P003 | Neutral |  |  | 0.11 | fixN | 4937 |  |
| P1929 | Neutral |  |  | 0.18 | fixN | 5236 |  |
| P1516 | Neutral |  |  | 0.05 | fixN | 5921 |  |
| P015 | Neutral |  |  | 0.15 | fixN | 6662 |  |
| P1798 | Neutral |  |  | 0.02 | typL | 7379 |  |
| P1464 | Neutral |  |  | 0.09 | fixN | 5553 |  |
| P1732 | Balancing |  |  | 0.08 | fixN | 1920 |  |
| P041 | Balancing |  |  | 0.14 | fixN | 2612 |  |
| P789 | Positive BayeScan |  |  | **0.25** | optN | 2807 |  |
| P1714 | Positive BayeScan |  |  | 0.03 | optB | 2880 |  |
| P1855 | Positive BayeScan |  |  | 0.15 | optN | 2971 |  |
| P1222 | Balancing Arlequin |  |  | 0.02 | fixN | 5647 |  |
| P414 | Balancing Arlequin |  |  | 0.13 | optN | 6309 |  |
| P550 | Balancing Arlequin |  |  | 0.06 | NullModel | na |  |
| P443 | Balancing Arlequin |  |  | 0.16 | optN | 2596 |  |

**Table S6.-** Summary of Mantel and Partial Mantel tests calculated conducting genetic distance (*F*_ST_) using the total set of loci, the putatively neutral dataset and the set of candidate loci to positive selection. Geographic distance (Geo) was calculated as the shortest water distance and temperature distance (Temp) refers to the average temperature measured in summer between 2005 and 2012 at 200 m depth. Partial Mantel tests are corrected by controlling for the effect of the matrix indicated in brackets. Boldface font depicts statistically significant *P*-values.

| **Test** | **Markers** | **Matrices** | **r_xy_** | ***P*-value** |
| --- | --- | --- | --- | --- |
| **Mantel Test** | All loci | Geo_*F*_ST_ | 0.784 | **0.0001** |
|  |  | Temp_*F*_ST_ | 0.712 | **0.0001** |
|  |  | Geo_Temp | 0.790 | **0.0001** |
|  | Neutrals | Geo_*F*_ST_ | 0.761 | **0.0001** |
|  |  | Temp_*F*_ST_ | 0.601 | **0.0001** |
|  | Positive selection | Geo_*F*_ST_ | 0.765 | **0.0001** |
|  |  | Temp_*F*_ST_ | 0.745 | **0.0001** |
| **Partial Mantel test** | All loci | Geo_*F*_ST_(Temp) | 0.514 | **0.0005** |
|  |  | Temp_*F*_ST_(Geo) | 0.243 | **0.0064** |
|  | Neutrals | Geo_*F*_ST_(Temp) | 0.584 | **0.0003** |
|  |  | Temp_*F*_ST_(Geo) | 0.000 | 0.9972 |
|  | Positive selection | Geo_*F*_ST_(Temp) | 0.430 | **0.0024** |
|  |  | Temp_*F*_ST_(Geo) | 0.358 | **0.0004** |

**FIGURES**

a)

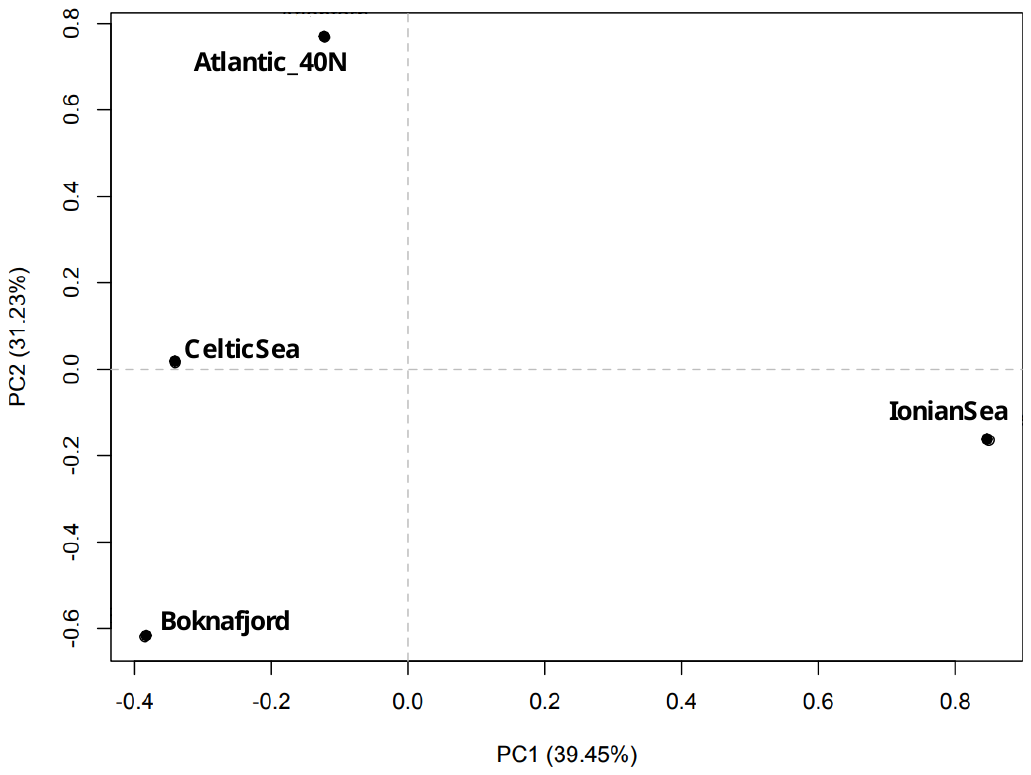

b)

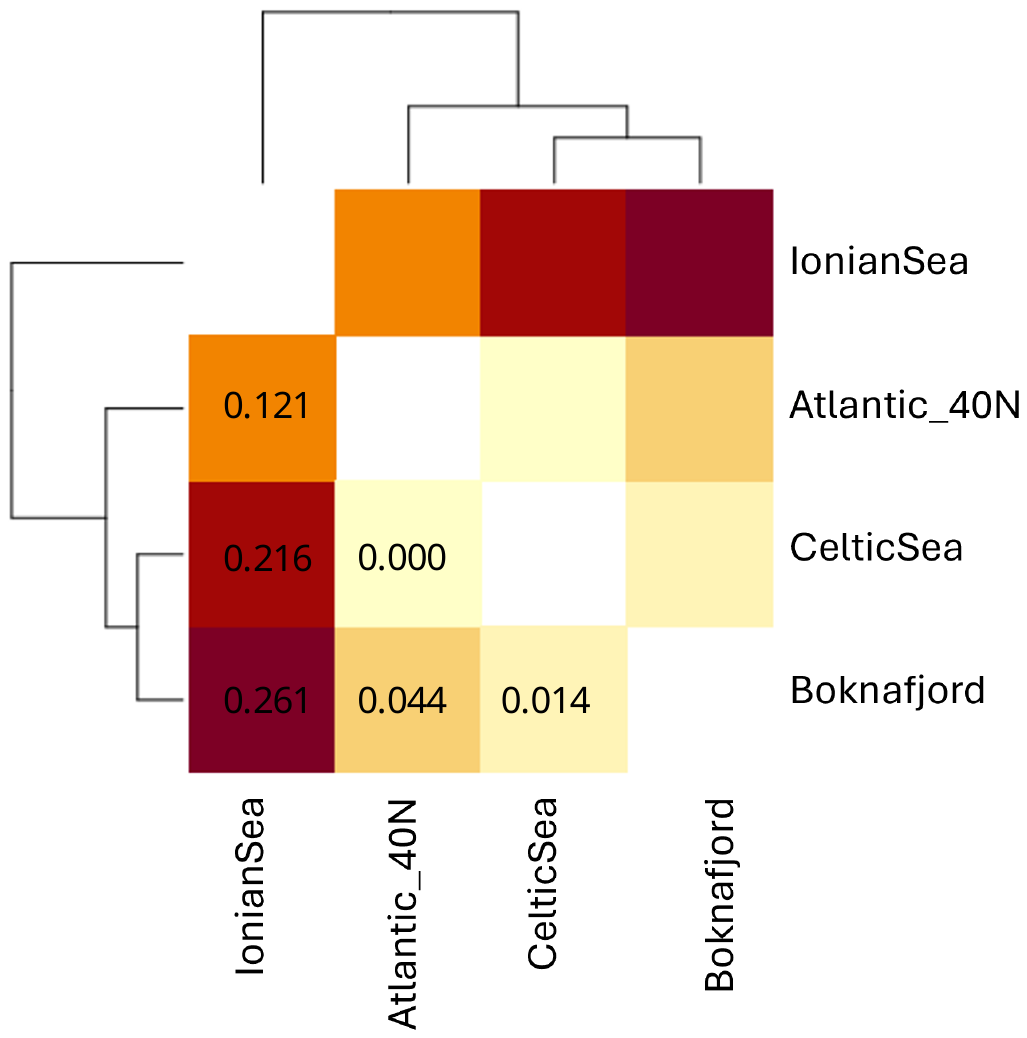

**Fig. S1.** Principal Component Analysis (a), and pairwise *F*_ST_ heatmap coupled with dendrogram built with the 12,000 filtered SNPs obtained from pool sequencing the four samples subsequently used to mine SNPs for throughput genotypying.

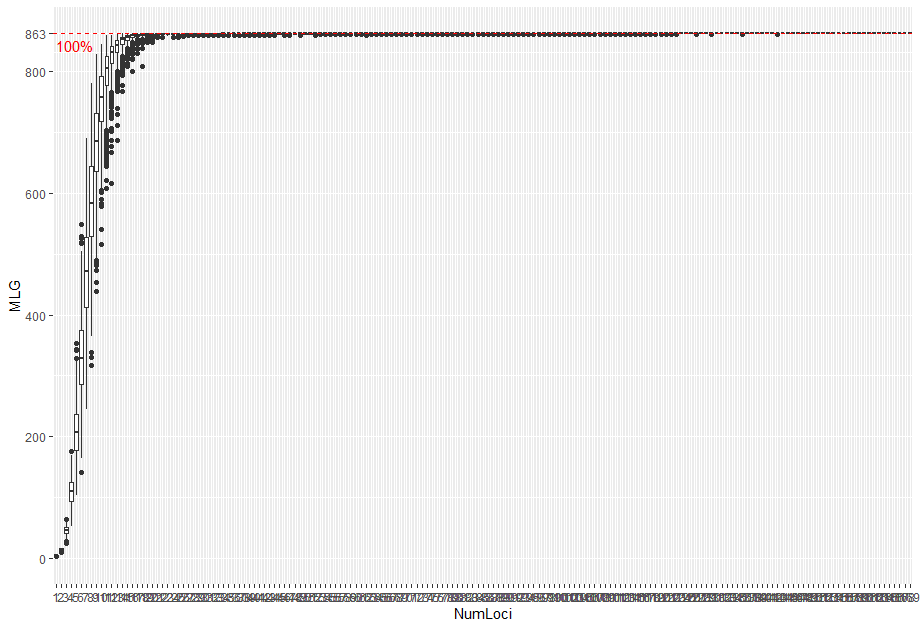

**Fig. S2.** Genotype accumulation curve calculated for the set of 170 polymorphic SNP loci using the total 863 individuals.

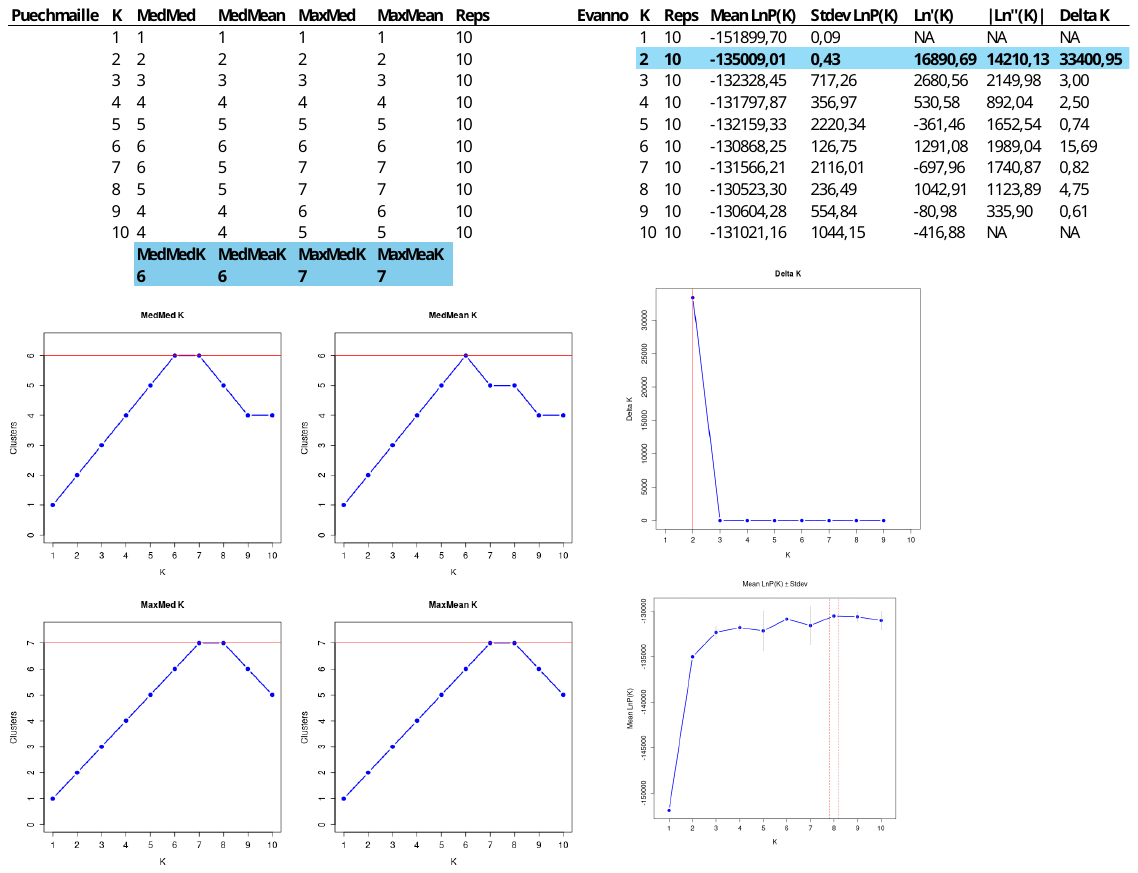

**Fig. S3.** *A posteriori* analysis of STRUCTURE outcome for the set of 170 loci following Puechmaille and Evanno’s statistics, respectively.

a)

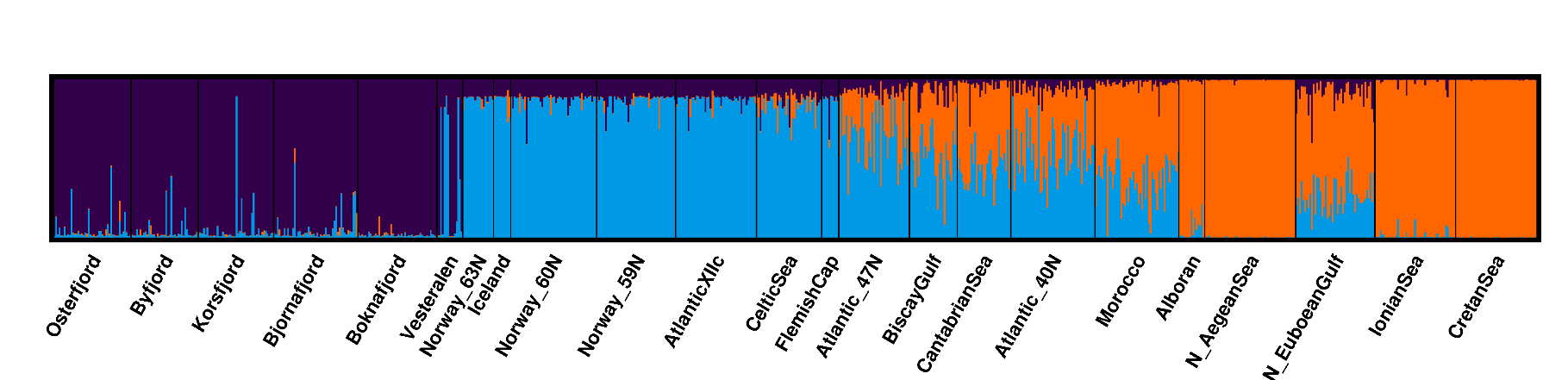

b)

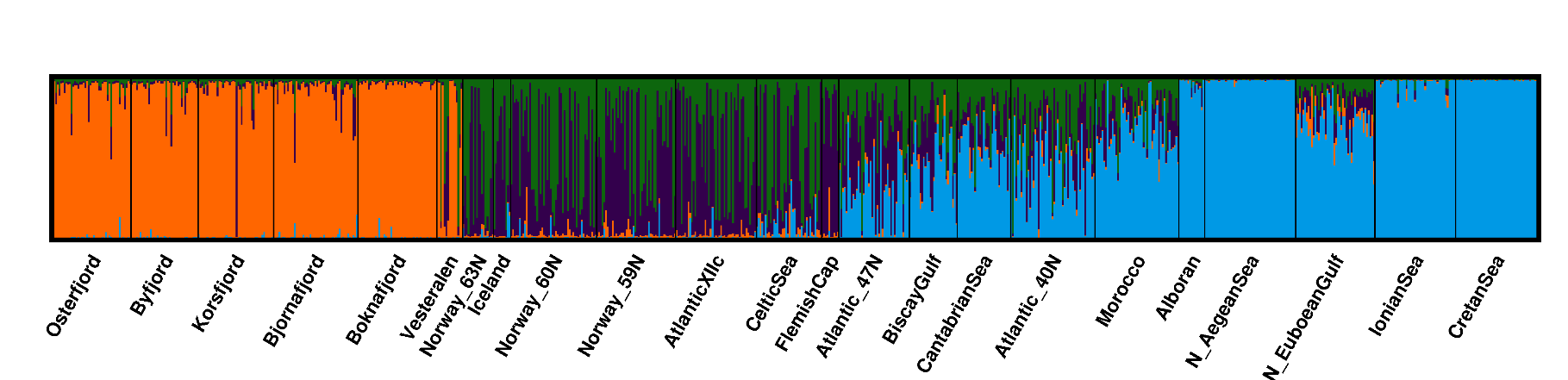

c)

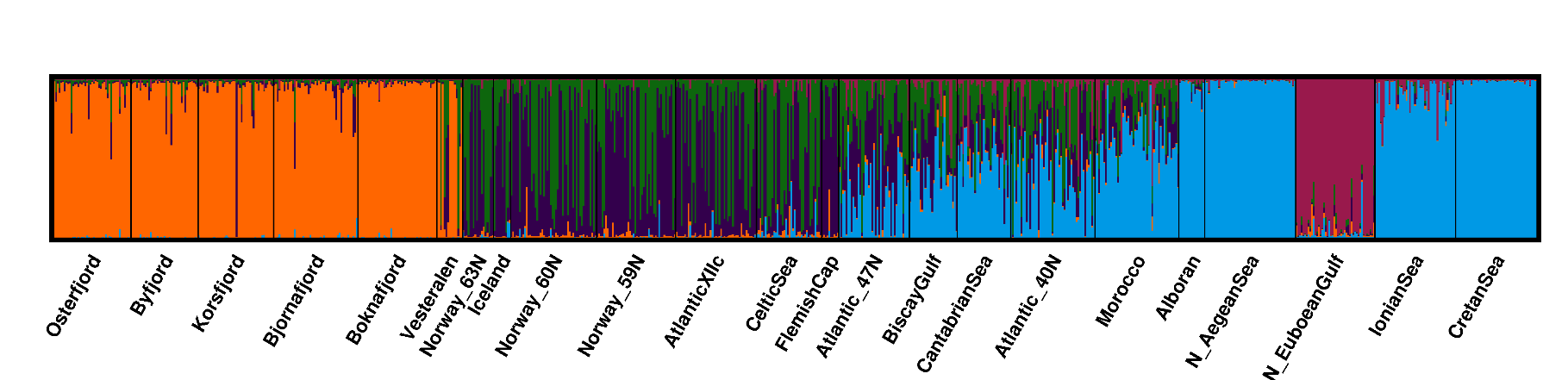

**Fig. S4.** Barplot representing the proportion of individuals’ ancestry to cluster at K3 to K5 after Bayesian clustering in STRUCTURE using the 170 total loci.

a)

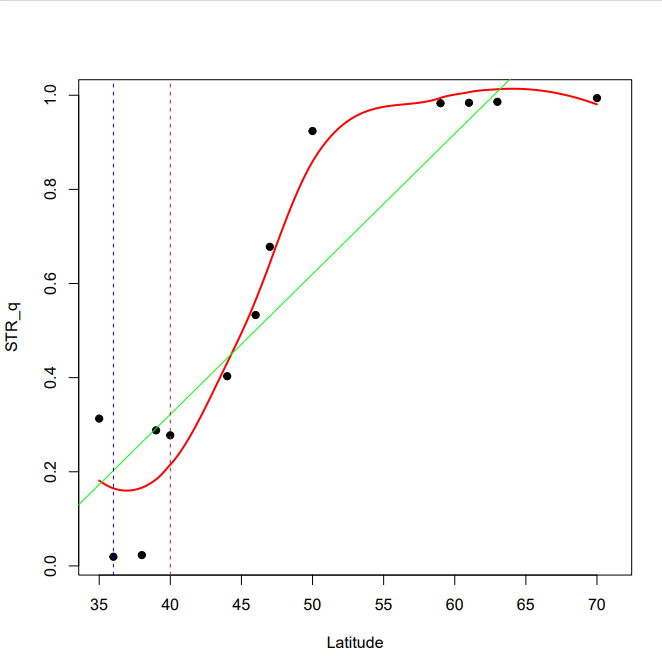

b)

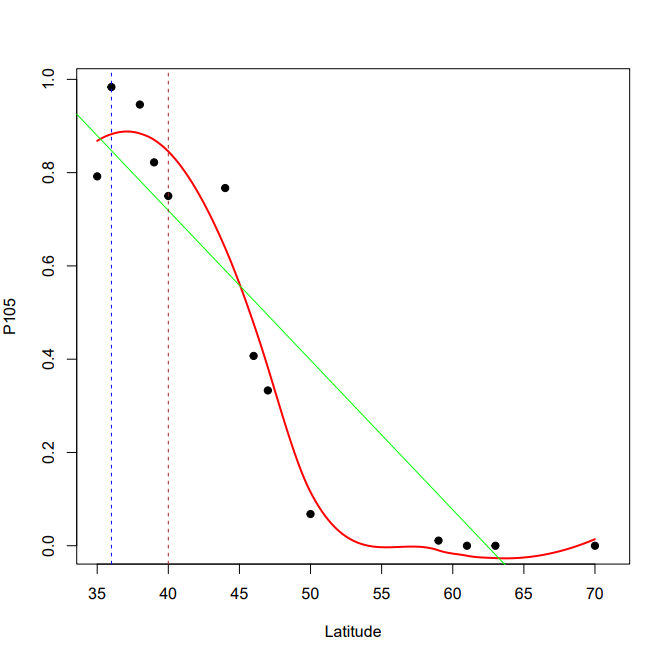

**Fig. S5.** Examples of cline latitudinal patterns for STRUCTURE Q-score (a) and locus P105 (b).

a)

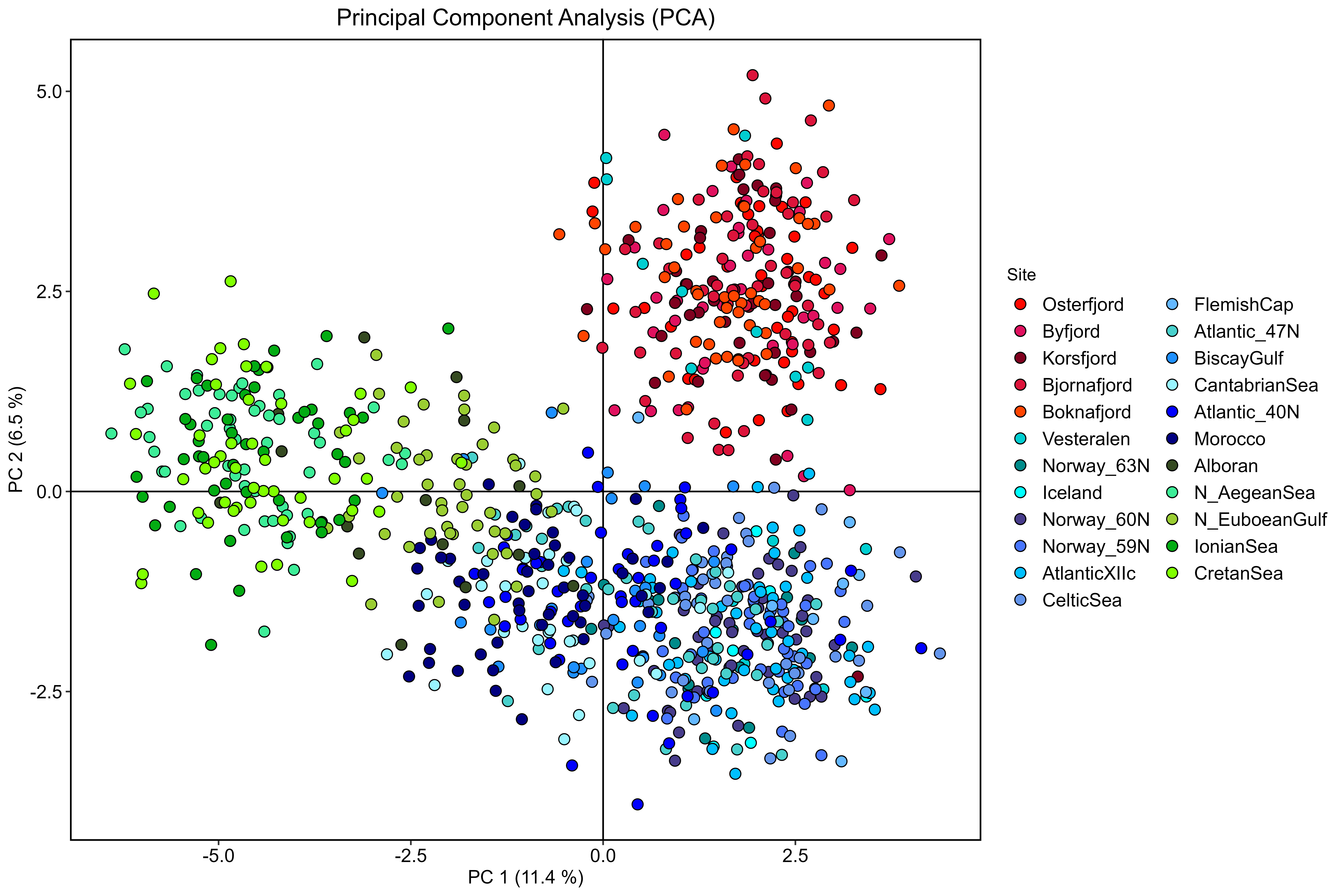

b)

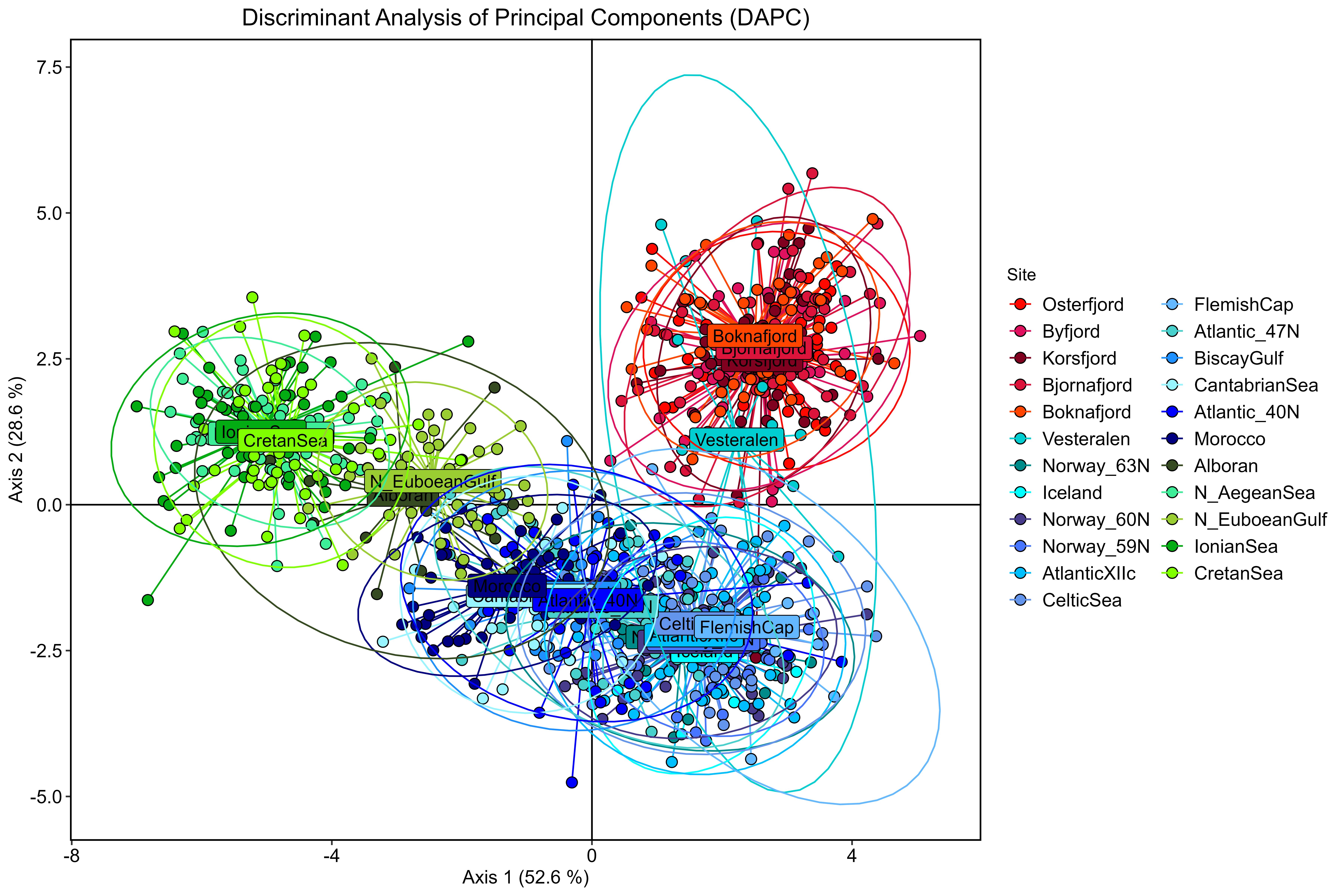

**Fig. S6.** Principal Component Analysis (PCA) of *Maurolicus muelleri* genotyped at 78 putatively neutral loci (a). Genetic differentiation using Discriminant Analysis of Principal Components (DAPC) after retaining 40 principal components and 3 discriminant functions (b). Blue dots depict the Ocean samples, red dots indicate fjord individuals and green dots represent the Mediterranean ones.

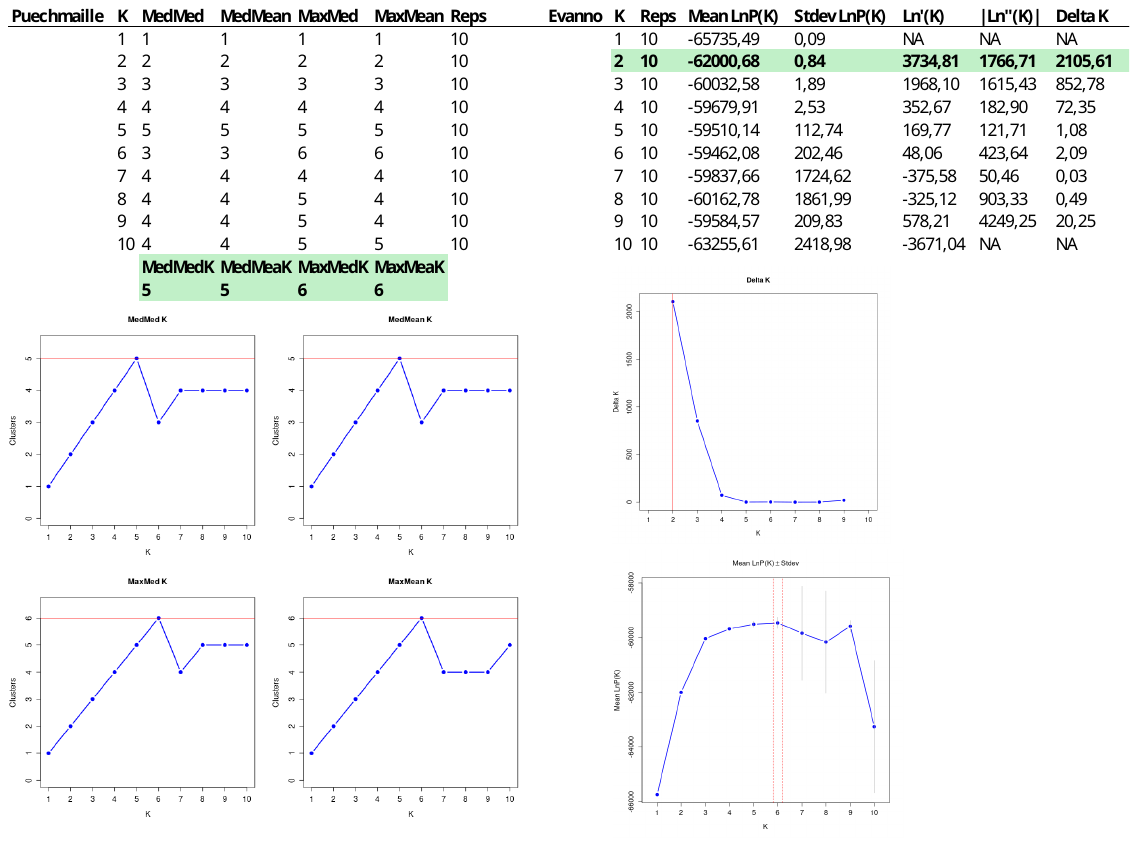

**Fig. S7.** *A posteriori* analysis of STRUCTURE outcome for the set of 78 loci following Puechmaille and Evanno’s statistics, respectively.

a)

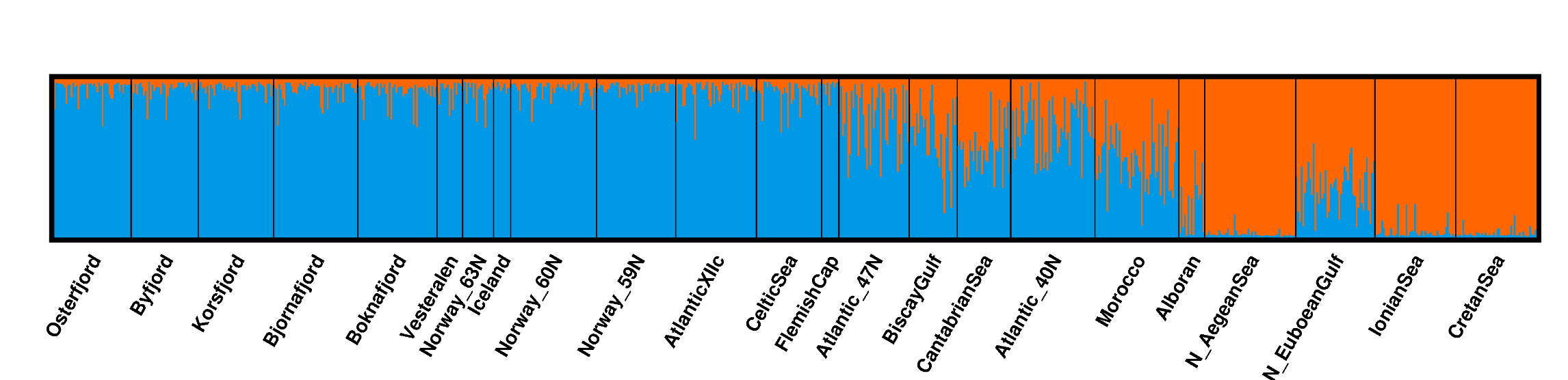

b)

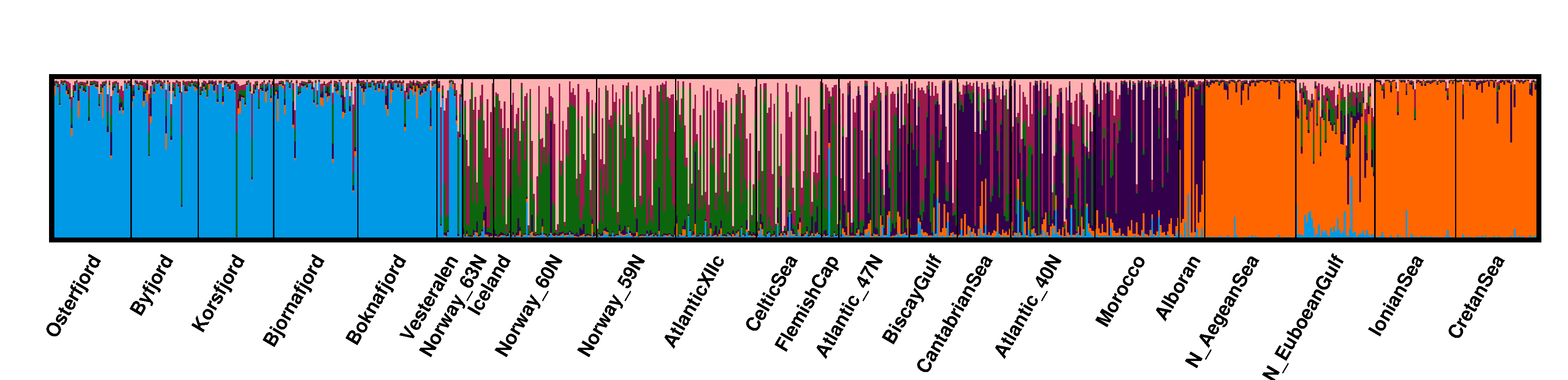

**Fig S8.** Barplot representing the proportion of individuals’ ancestry to cluster at a) K2 and b) K6 after Bayesian clustering in STRUCTURE using the 78 putatively neutral loci as determined by the *a posteriori* analyses conducted using Evanno’s test and Puechmaille’s statistics, respectively.

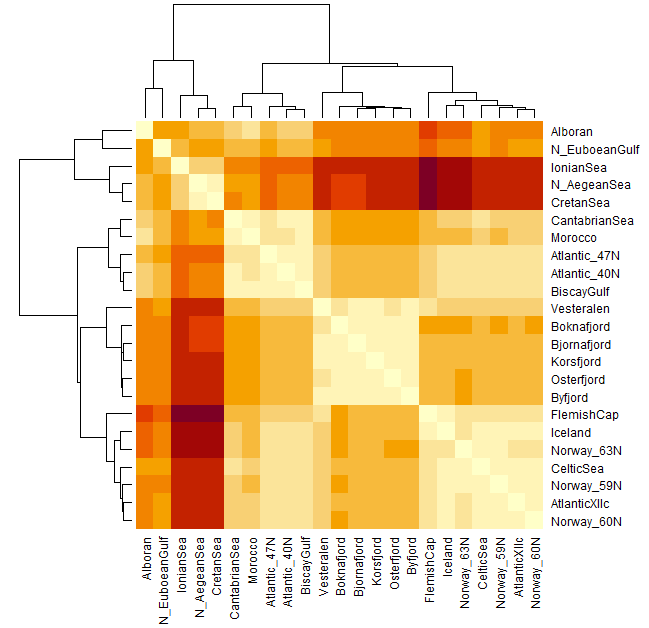

**Fig S9.** Heatmap *F*_ST_ coupled with dendrogram for the set of 78 putatively neutral markers*.* Pairwise values and significance can be found in Suppl. Table S3. Colours depict the degree of differentiation: beige colours indicating lower differentiation and moving towards dark brown to indicate larger differentiation.

a)

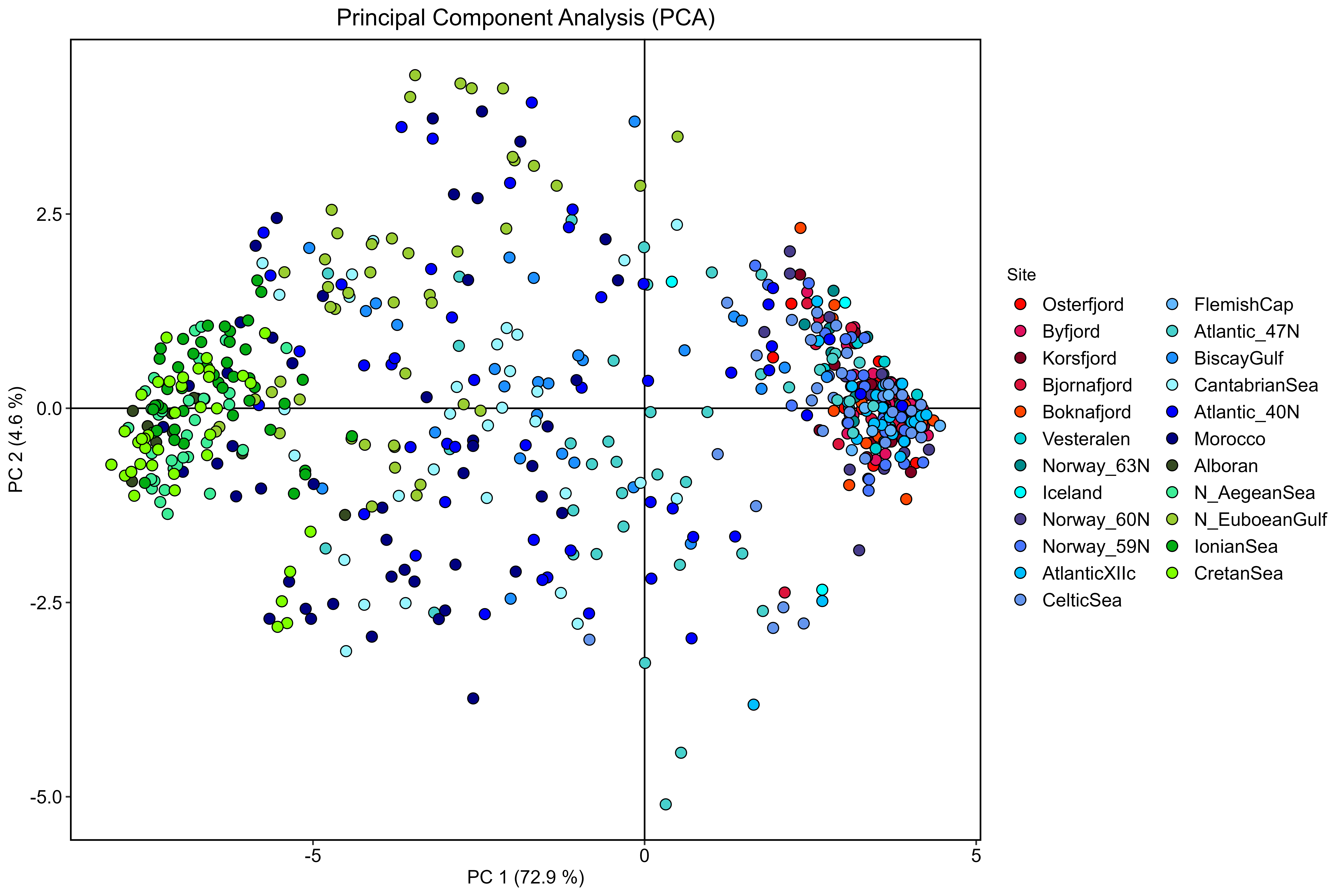

b)

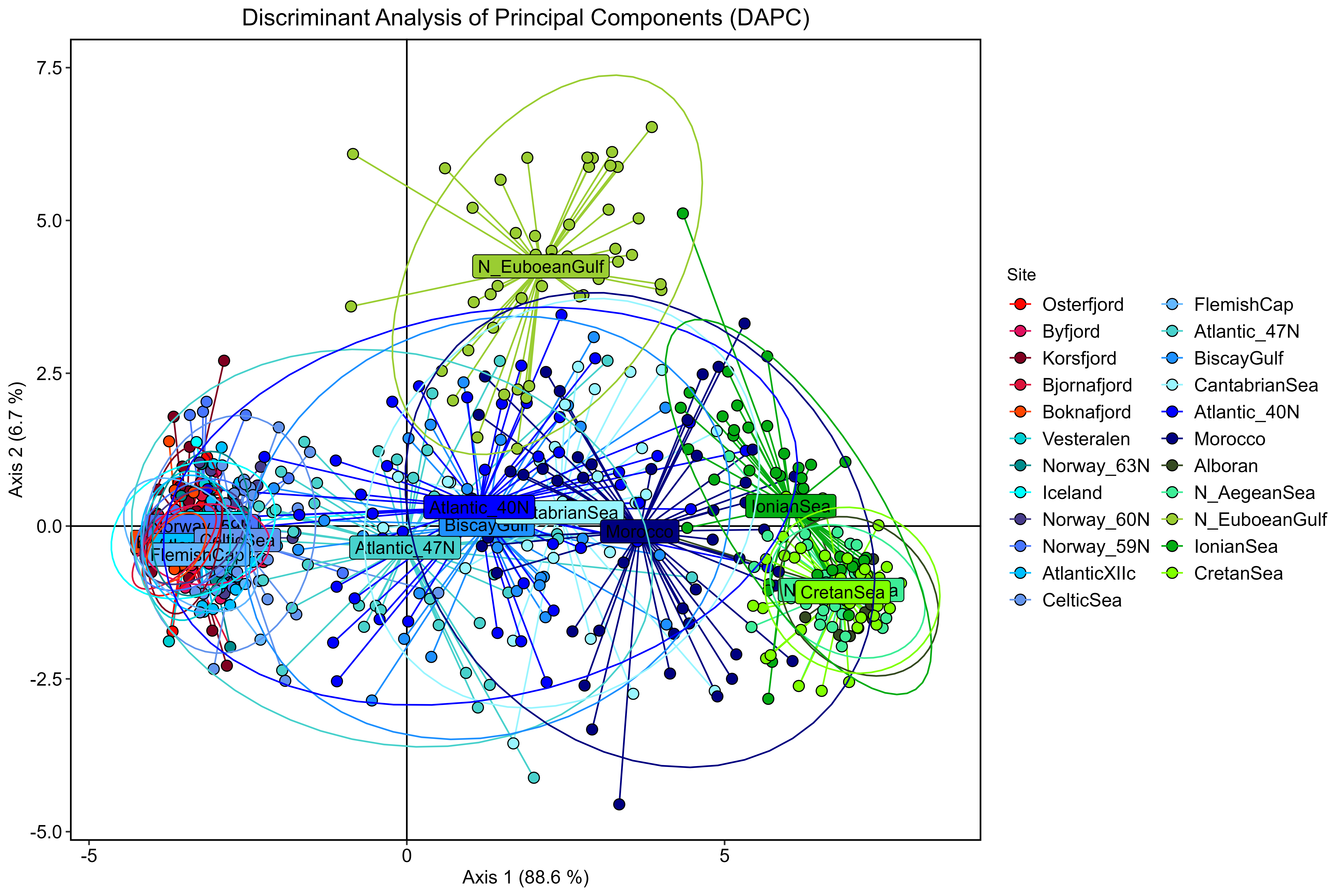

**Fig. S10.** Principal Component Analysis (PCA) of *Maurolicus muelleri* genotyped at 20 candidate outliers to positive selection (a). Genetic differentiation using Discriminant Analysis of Principal Components (DAPC) after retaining 18 principal components and 3 discriminant functions (b). Blue dots depict the Ocean samples, red dots indicate fjord individuals and green dots represent the Mediterranean ones.

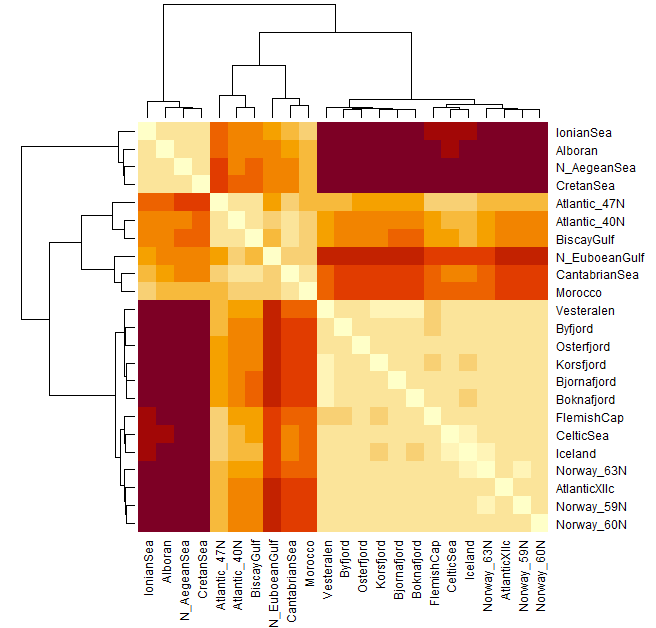

**Fig S11.** Heatmap *F*_ST_ coupled with dendrogram for the set of 20 candidate loci to positive selection*.* Pairwise values and significance can be found in Suppl. Table S4. Colours depict the degree of differentiation: beige colours indicating lower differentiation and moving towards dark brown to indicate larger differentiation.

a)

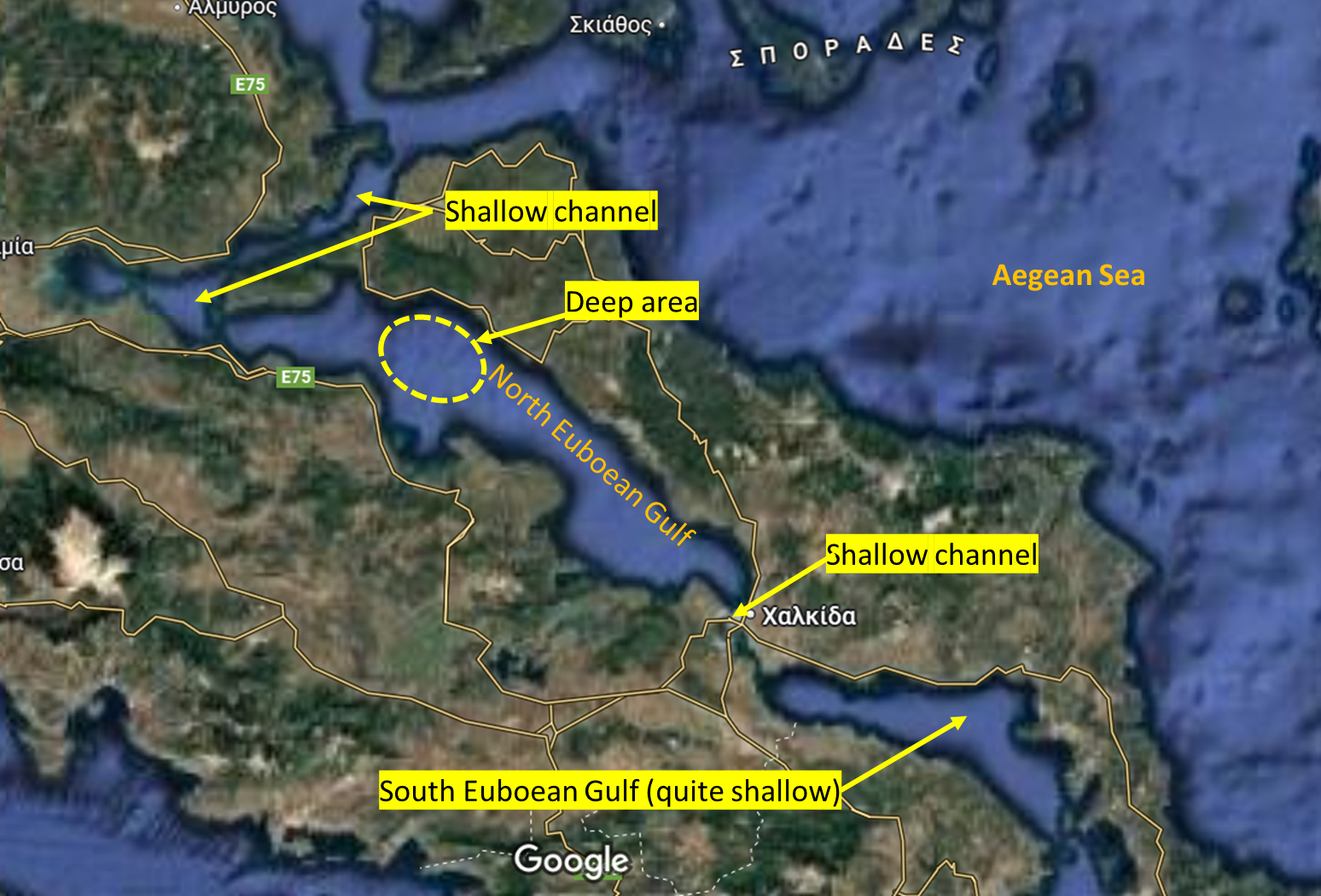

b)

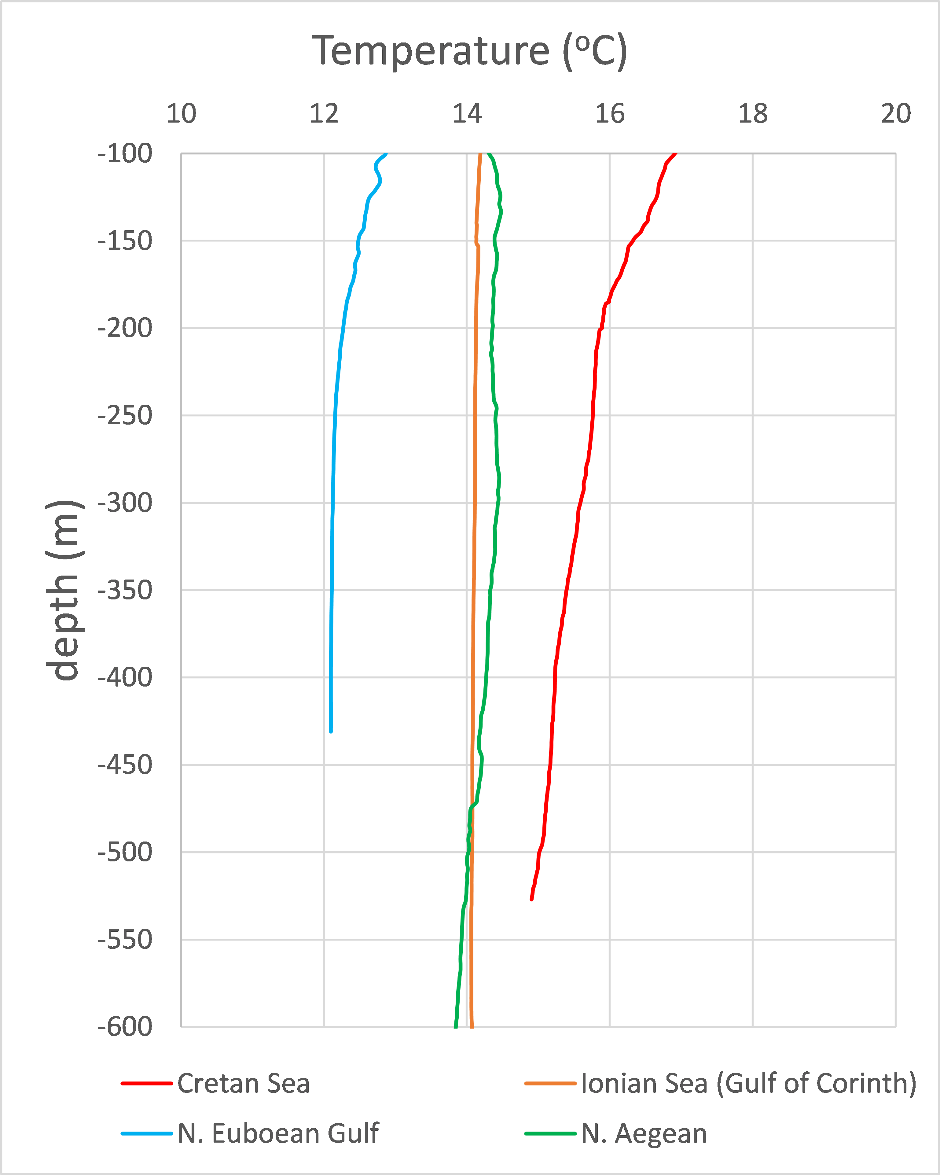

**Fig S12.** Detailed map of the N. Euboean Gulf (a) and temperature at depth measured with CTD for the sites sampled in the Greek seas (b).
